# RNAP-seq: *in vitro* genome-scale transcription reveals preferential RNA polymerase pausing on *Clostridioides difficile* antisense DNA

**DOI:** 10.64898/2026.05.06.723315

**Authors:** Ping Shen, Michael B. Wolfe, Jason Saba, Rachel Mooney, Yu Bao, Ezaz Ahmad, Robert Landick, Xinyun Cao

**Affiliations:** Department of Microbiology, University of Texas Southwestern Medical Center, Dallas, TX, 75390, USA; Department of Biochemistry, University of Wisconsin-Madison, Madison, WI, 53706, USA; Department of Bacteriology, University of Wisconsin-Madison, Madison, WI, 53706, USA

## Abstract

Mechanistic dissection of gene transcription and its regulation in diverse organisms remains a central challenge for molecular biology. *In vivo* analyses using high-throughput DNA sequencing readouts (e.g., RNA-seq, Term-seq, and NET-seq) have revealed that transcriptional pausing by RNA polymerase (RNAP) underpins the regulation of gene transcription. However, these *in vivo* methods are difficult to apply in non-model organisms and lack the power of bottom-up biochemical analysis that can deconvolute the confounding effects of the cellular environment. Here we report a method for unbiased transcription of entire genomes *in vitro* called RNAP-seq, which is capable of defining pause, arrest, and termination sites with single-nucleotide resolution and of defining the regulatory contributions of *cis*-acting RNA and DNA sequences and dissociable transcription factors with complete control over transcription conditions. Using RNAP-seq to compare transcription by recombinant RNAPs from *Clostridioides difficile* (*Cdf*) and *Escherichia coli* (*Eco*), we discovered that *Cdf*RNAP exhibits pausing signatures distinct from *Eco*RNAP. *Cdf*RNAP pauses more frequently and more strongly at T-rich sequences, particularly on antisense regions of genes. We also defined the action of *Cdf*NusG at genome scale, revealing its preferential action in riboswitch control regions. These findings demonstrate lineage-specific pausing features and suggest that genome composition and RNAP co-evolve to shape gene expression. RNAP-seq is broadly adaptable to diverse bacterial species and offers a powerful framework for uncovering fundamental principles of transcription that govern gene expression.

**Highlights:**

- RNAP-seq enables unbiased transcription of every base-pair in an organism’s genome
- RNAP-seq reveals distinct *E. coli* (*Eco*) and *C. difficile* (*Cdf*) pause sequences
- *Cdf*NusG broadly enhances pausing and termination, notably in riboswitch-encoding DNA
- *Cdf*RNAP but not *Eco*RNAP pauses more on the T-rich antisense strand of *Cdf* genes

## INTRODUCTION

The growth and survival of all living organisms rely on precise control of gene expression. RNA polymerase (RNAP), a multi-subunit protein machine that transcribes genetic information from DNA into RNA, is the central catalytic and regulatory platform for this control. Transcription proceeds through three major stages: initiation, elongation, and termination. Notably, elongation is highly regulated. This process of RNA synthesis is not continuous but is frequently interrupted by transcriptional pausing, which occurs on average once every ∼100 base pairs *in vivo* in the model organism *Escherichia coli*^1^. During these pauses, RNAP remains bound to the nucleic acid while temporarily ceasing nucleotide addition. Pausing by RNAP is a fundamental regulatory feature conserved across prokaryotes and eukaryotes^2–6^.

In bacteria, pausing synchronizes transcription with translation and creates regulatory windows for key molecular events, including metabolite binding to riboswitches, recruitment of transcription factors (TFs), proper folding of nascent RNA, and the opportunity for transcription termination^4^. Despite its importance, our understanding of RNAP pausing at the genome scale is mainly derived from *E. coli*. However, the mechanisms governing RNAP pausing and its regulation are likely to vary across phylogenetically diverse bacterial lineages. For example, NusG, the only universally conserved transcription factor, inhibits pausing in *E. coli*^7,8^ but stimulates pausing in *Bacillus subtilis*^9^ and *Mycobacterium tuberculosis*^10^. Differences also may arise from intrinsic RNAP properties (*e.g.* lineage-specific insertions that alter enzyme kinetics), and genome composition, as bacteria span a wide range of AT- to GC-rich sequences, and TFs that exhibit divergent functions across species.

Genome-wide mapping of transcription elongation has been enabled by *in vivo* approaches such as Term-seq and native elongating transcript sequencing (NET-seq)^11,12^. Although powerful, these methods have important limitations. Pausing *in vivo* is shaped by a complex network of factors including DNA topology, R-loop formation, translational coupling, TFs (e.g., NusA and NusG), small-molecule regulators, and nucleoid-associated proteins^4^. The complexity of this network combined with the fact that transcription is essential for cell viability make it difficult to isolate the contributions of individual components of RNAP action. NET-seq also requires chromosomal tagging of RNAP for immunoprecipitation, which is challenging for unculturable or genetically intractable organisms. To date, nearly all *in vitro* studies of transcript elongation that provide a controlled environment for mechanistic studies have been limited to single fragments of DNA less than 1 kb in length. Genome-scale methods for *in vitro* transcription have been devised but existing approaches have been primarily limited to use of microarrays or 5′ RNA end mapping to study the regulation of transcription initiation^13–17^. Recently, 3′-end mapping was reported for *in vitro* genome-scale transcription using *M. tuberculosis* RNAP but only termination sites were detected and most of the genome was not analyzed because transcription relied on native promoters with limited activity *in vitro*^13^. Thus, a method to assay RNAP pausing at genome scale under defined biochemically manipulable conditions remains lacking and highly desirable.

To fill this gap, we have developed a robust *in vitro* transcription system termed RNAP-seq (<u>RNA</u> <u>p</u>olymerase <u>seq</u>uencing) that assays RNAP pausing genome-wide at single-nucleotide resolution (Figure 1A). This highly controllable approach relies on purified recombinant proteins and eliminates the confounding complexity of the cellular environment, enabling precise dissection of pausing regulation by both cis-acting sequence elements and accessory transcription factors (TFs). Leveraging this system, we directly compared *E. coli* (*Eco*) RNAP to *Clostridioides difficile (Cdf)* RNAP. *Cdf* is a deadly human pathogen distantly related to *Eco* with a markedly different AT-rich genome (∼70% AT versus ∼50% in *Eco*). Its transcriptional pausing mechanisms are unexplored. RNAP-seq allowed comparison of *Eco*RNAP and *Cdf*RNAP using both cognate and heterologous genomic templates (Figure 1B). This design allows us to pinpoint the relative contributions of RNAP intrinsic properties and genome sequence context to transcriptional pausing.

**Figure 1:**
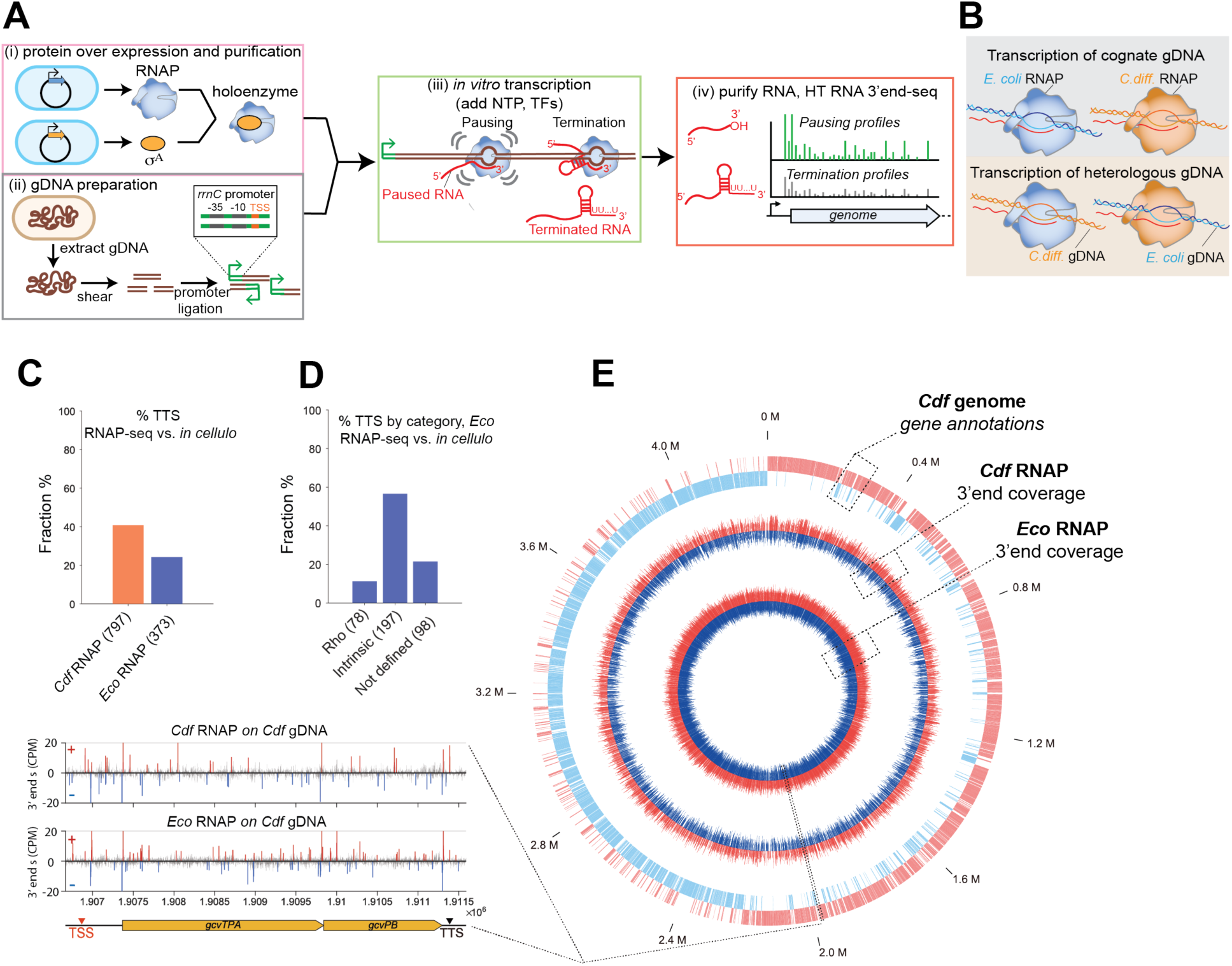
Schematic workflow and scope of RNAP-seq. **A**, Schematic of the RNAP-seq workflow. RNAP-derived 3′ ends include signals arising from both transcriptional pausing and termination. Transcription termination sites (TTS) are identified and excluded from the full set of 3′ ends using a bioinformatic pipeline (Materials and Methods). **B**, Schematic illustrating RNAP-seq transcription of cognate genomic DNA (gDNA, top) and heterologous genomic DNA (bottom) by *Eco* and *Cdf* RNAP. **C**, Bar graph showing the fraction of TTS identified by RNAP-seq using *Cdf* RNAP on *Cdf* gDNA (orange) and *Eco* RNAP on *Eco* gDNA (blue), relative to *in cellulo* TTS datasets^18,19^. If multiple termination sites identified by RNAP-seq fell within a ±5 nt window of a terminator in the corresponding in cellulo dataset, they were considered a single site. The total number of TTS detected by RNAP-seq for *Cdf* and *Eco* RNAP is indicated on the X-axis. **D**, Bar graph showing the fraction of intrinsic, Rho-dependent, and unclassified TTS identified by RNAP-seq using *Eco* RNAP on *Eco* gDNA, relative to the *in cellulo* dataset^18^. The number of TTS detected by RNAP-seq in each category is indicated on the X-axis. **E,** Genome wide distribution of 3′ end halt signals on the *Cdf* genome. Outer circle: gene annotations; middle circle: 3′ end coverage by *Cdf* RNAP; inner circle: 3′ end coverage by *Eco* RNAP. Red and blue colors represent positive and negative strands, respectively. Mb, megabase.

## RESULTS

### RNAP-seq enables unbiased transcription of every base pair in an organism’s genome

To enable uniform and efficient transcription across an entire genome, we ligated a strong *Cdf* ribosomal RNA (*rrnC*) promoter to sheared genomic DNA (gDNA) from *Cdf* or *Eco* (Figure 1A). This approach bypasses the need for RNAP to initiate at native promoters, which often require specific transcriptional activators and may have little or no initiation activity *in vitro*. Use of a strong promoter appended to gDNA fragments ensures robust RNA output for deep sequencing coverage of transcript 3′ ends. The *Cdf rrnC* promoter is also recognized by *Eco*RNAP, which makes RNAP-seq easily adaptable to broad bacterial species.

Transcription was initiated by addition of equimolar NTPs (1 mM each) to RNAP–χρ holoenzyme mixed with the promoter–gDNA fragments. As nascent RNA was produced, RNAP released χρ–promoter contacts and encountered sites in the gDNA that govern RNAP pausing and termination (Figure 1A). Attachment of the *rrnC* promoter significantly increased transcript yield compared to non-ligated controls. For *Eco*RNAP, the total transcript yield increased 8-fold compared to gDNA without an attached promoter (Figure S1A). Total RNAs generated from each reaction were ligated to 3′-adapter and used to generate Illumina-compatible cDNA libraries for high-throughput sequencing (Figure S1B). Each sequencing read primed from the 3′-adapter identifies the 3′ end of a transcript, which enabled single nucleotide–resolution mapping of RNAP activity across the gDNA.

To characterize RNAP transcription patterns, we developed a custom bioinformatic pipeline that removes background noise and extreme high-signal outliers. To eliminate outlier signals, we first identified transcription termination sites (TTS) from *in vivo* datasets for both *Eco*^18^ and *Cdf*^19^ that are also detected by RNAP-seq (Figures 1C and 1D). We then used the read distributions of these termination signals to establish lower and upper bounds for signal filtering. We identified high-confidence transcription halt sites by iterative peak-calling based on local enrichment (*z*-score) and signal intensity^1,20^. In each iteration, positions with z-score >4 within a 200 bp window were classified as halt sites (Figure S1C, and Materials and Methods). These high-confidence halt sites represent genomic positions at which RNAP accumulates (due to pausing or arrest) or releases transcripts (due to intrinsic termination) (Figure S1D). We refer to these positions as RNAP halt sites (resulting from pausing, arrest, or termination), and the associated signal as an RNAP halt signal to avoid implying a specific mechanism prior to characterization.

The genome-wide distribution of RNAP halt sites is relatively uniform across both *Eco* and *Cdf* genomes (Figures. 1E, S2A). In total, we identified 11,410 halt sites for *Eco*RNAP on *Eco* gDNA, 16,565 for *Eco*RNAP on *Cdf* gDNA, 12,533 for *Cdf*RNAP on *Cdf* gDNA, and 3,908 for *Cdf*RNAP on *Eco g*DNA (Figure S2B, Supplementary Tables 1-4), representing sites consistently detected across all three replicates. To confirm that these signals arise from RNAP activity rather gDNA fragmentation, we compared RNAP-seq signals with sequencing profiles of the gDNA fragment ends (Figure S3). RNAP-seq derived halt sites showed no enrichment at sheared DNA ends, indicating that the detected peaks reflect bona fide transcriptional events rather than library preparation artifacts.

To quantify signal strength, we defined a RNAP halt score as the read count at each halt site normalized to the mean read count within a 200-nt window centered on that site (±100 nt). This normalization accounts for local background transcription and enables comparison across genomic contexts. Importantly, the halt score reflects the relative accumulation of RNAP at a given position rather than being a direct measure of pausing kinetics. Halt scores were highly reproducible between biological replicates (Figure. S4), supporting the robustness of the method. We conclude that RNAP-seq provides a robust, quantitative and genome-wide readout of RNAP halt sites, capturing transcriptional pausing, arrest, and termination events.

### RNAP-seq captures genome-wide transcription termination and kinetically stable RNAP pause states, separable by a bioinformatic pipeline

We reasoned that RNAP-seq detects three major classes of RNA 3′ ends: (1) pausing-derived 3′ ends, which correspond to paused elongation complexes (PECs)^4^; (2) arrested RNAP states, including halted or catalytically inactive complexes that fail to resume elongation^21,22^; and (3) intrinsic-termination 3′ ends generated by terminator hairpin formation at PECs with poly-U RNA–DNA hybrids, leading to transcript release^23^ (Figure S1D). PECs form at multiple types of pause sites, including elemental pauses (ePECs), backtracked pauses, and long-lived (*i.e.* kinetically stable) pauses^4^. These pause events are all reversible states from which RNAP can resume transcription, in contrast to arrest sites at which RNAP becomes irreversibly stuck. Arrested or long-lived pause events are likely to be more common during RNAP-seq than during *in vivo* transcription because RNAP-seq integrates RNAP activity over extended *in vitro* transcription reaction times (15 min incubation) in the absence of RNAP-rescue and other TFs and during which time PPi accumulates. As a result, events that may be transient *in vivo* accumulate into an extensive background while kinetically stable pauses and termination events become preferentially enriched.

To distinguish transcription termination sites from pausing and arrest sites, we first analyzed *Eco*RNAP transcribing *Eco* gDNA. RNA 3′ ends that fell within a ±5 nt window of intrinsic terminators defined from an *in vivo* dataset^18^ were classified as terminator-associated. Sequence logo analysis of these sites revealed canonical intrinsic terminator features, including a downstream poly-T tract and an upstream GC-rich region corresponding to a terminator RNA hairpin (Figures 2A, 2B). This assignment was further supported by RNA secondary structure predictions, which showed that these sites are associated with more stable predicted upstream hairpins than either random sites or enriched halt sites not associated with terminators (*i.e.*, lower ΔG) (Figure 2C). Based on these analyses, we established a bioinformatic pipeline to systematically separate terminator-associated sites from pausing and arrest sites (Figure S1E).

**Figure 2:**
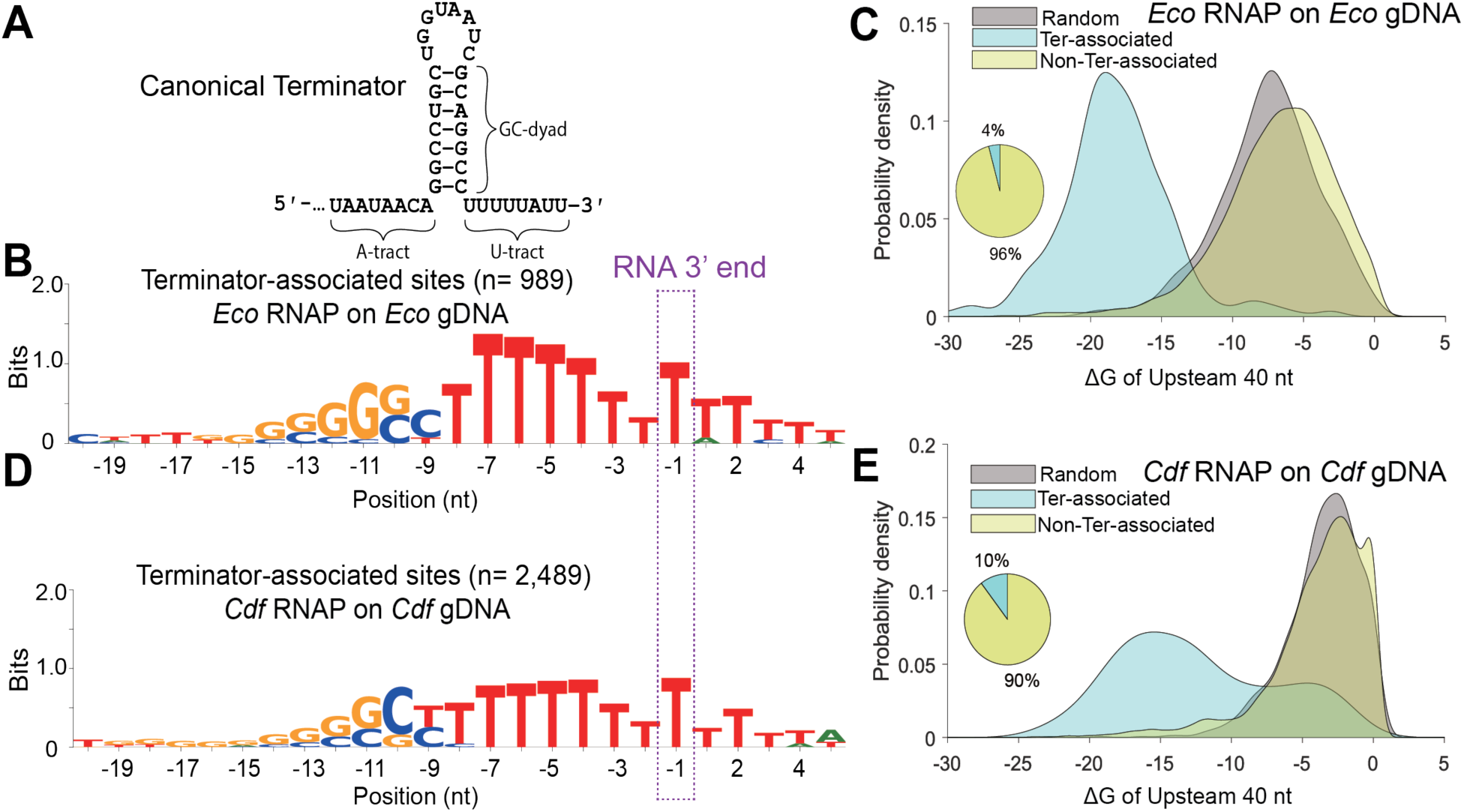
Sequence determinants underlying intrinsic terminations and transcriptional pausing. A, Sequence and secondary structure of a canonical λtR_2_ intrinsic terminator **B**, 3′ end halt sites were classified as terminator-associated (“Ter-related”) when located upstream of *in cellulo* intrinsic terminators. Sequence logos show nucleotide frequencies surrounding the RNA 3′ end (position -1). Ter-related 3′ end display a characteristic T-rich tract upstream of the 3′ end (-1), consistent with the U-tract typical of intrinsic transcription terminators, whereas non–terminator-associated sites show weaker sequence bias of poly-T. **C**, Distribution of predicted RNA hairpin stability upstream of RNA 3′ end (-1) on the *Eco* genome. Hairpin free energy (ΔG) was calculated for sequences located within 40 bp upstream of each 3′ end. Ter-related 3′ ends are associated with significantly more stable predicted RNA secondary structures (more negative ΔG) compared with non–terminator-associated 3′ ends or random genomic regions. The inset pie chart shows the fraction of corresponding 3′ ends populations. **D**, Sequence logos of 3′ ends generated by *Cdf* RNAP on the *Cdf* genome. **E**, Distribution of predicted RNA hairpin stability upstream of RNA 3′ ends on the *Cdf* genome.

Using a similar approach, we identified terminator-associated sites for *Cdf*RNAP on *Cdf* gDNA based on an *in vivo* Term-seq dataset (Figure 2D)^19^. We then could classify the remaining *Cdf*RNAP halt sites that were not associated with intrinsic terminators. RNA secondary structural analyses showed that these remaining halt sites lacked strong terminator-like sequence features and were associated with significantly less stable upstream RNA structures (*i.e.* higher predicted ΔG), resembling random genomic regions. In contrast, the terminator-associated sites were linked to more stable predicted upstream RNA hairpins (lower predicted ΔG values, Figure 2E).

Together, these results for *Eco*RNAP and *Cdf*RNAP indicate that our RNAP-seq analysis pipeline effectively separates transcription termination sites from non-terminator-associated halt sites, which thus predominantly reflect paused and arrested RNAP states.

### RNAP-seq captures elemental pausing by diverse RNAPs with distinct patterns from *in vivo* **NET-seq**

After removing intrinsic termination sites, we next analyzed the remaining non-terminator-associated halt sites (Figure 2C and 2E) to identify pausing patterns. Previous NET-seq studies in *E. coli* identified a consensus elemental pause (ce-pause) signal characterized by the sequence motif 5′ -g–11G–10t–3g–2Y−1G+1 (where –1 marks the 3′ end of the nascent RNA)^1^. Multiple sequence elements (*i.e.,* upstream and downstream fork junctions, the DNA–RNA hybrid, and downstream DNA) contribute additively to elemental pause strength^24^. Although this motif as first defined in *E. coli* causes pausing from bacterial to human RNAPs^1^, identifying the actual consensus pausing sequence in other bacteria by NET-seq has been challenging in part due to the difficulty of capturing RNAP at pause sites during cell recovery and lysis^9,20^. For some organisms, no consensus was detected^9^, and in others pauses appeared to occur either just before instead of after the consensus –1Y or one nucleotide further downstream from the –1 position^12,20,25,26^. The shifts in register may reflect RNAP reverse or forward translocation and transcript cleavage or nucleotide addition during sample processing^20^. Thus, the sequence determinants of elemental pausing in diverse organismal lineages has remained unclear.

To ask directly if *Cdf*RNAP and *Eco*RNAP recognize pause signals with the same or different consensus sequences, we applied RNAP-seq to each RNAP transcribing their respective gDNA templates. Strikingly, both RNAPs produced similar consensus pause sequences that resembled the ce-pause sequence obtained from NET-seq on *E. coli* (Figure 3)^1,20^. Both RNAPs preferentially slowed or halted at sites with prominent –11G/C, –10G/C, and –1Y features. A key difference from the sequence identified by NET-seq was observed at the +1 position for both RNAPs, wherein RNAP-seq consistently favored +1A rather than +1G observed in the *E. coli* NET-seq consensus. This bias likely reflects the equimolar NTP concentrations (1 mM each) used in RNAP-seq, whereas *in vivo* GTP is likely depleted during cell processing for NET-seq, exaggerating pauses before G. These observations are consistent with biochemical studies showing that both +1A and +1G strongly contribute to elemental pausing ^20,24^. Notably, prior *in vivo* NET-seq revealed that *Bacillus subtilis* RNAP exhibits a preference for +1A, consistent with our findings^1^. Despite these shared features, lineage-specific differences were also apparent. *Cdf*RNAP exhibited enrichment of T-rich sequences within the −9 to −3 region, whereas *Eco*RNAP showed enrichment of A-rich sequences in the same region, consistent across both RNAP-seq and NET-seq datasets.

**Figure 3:**
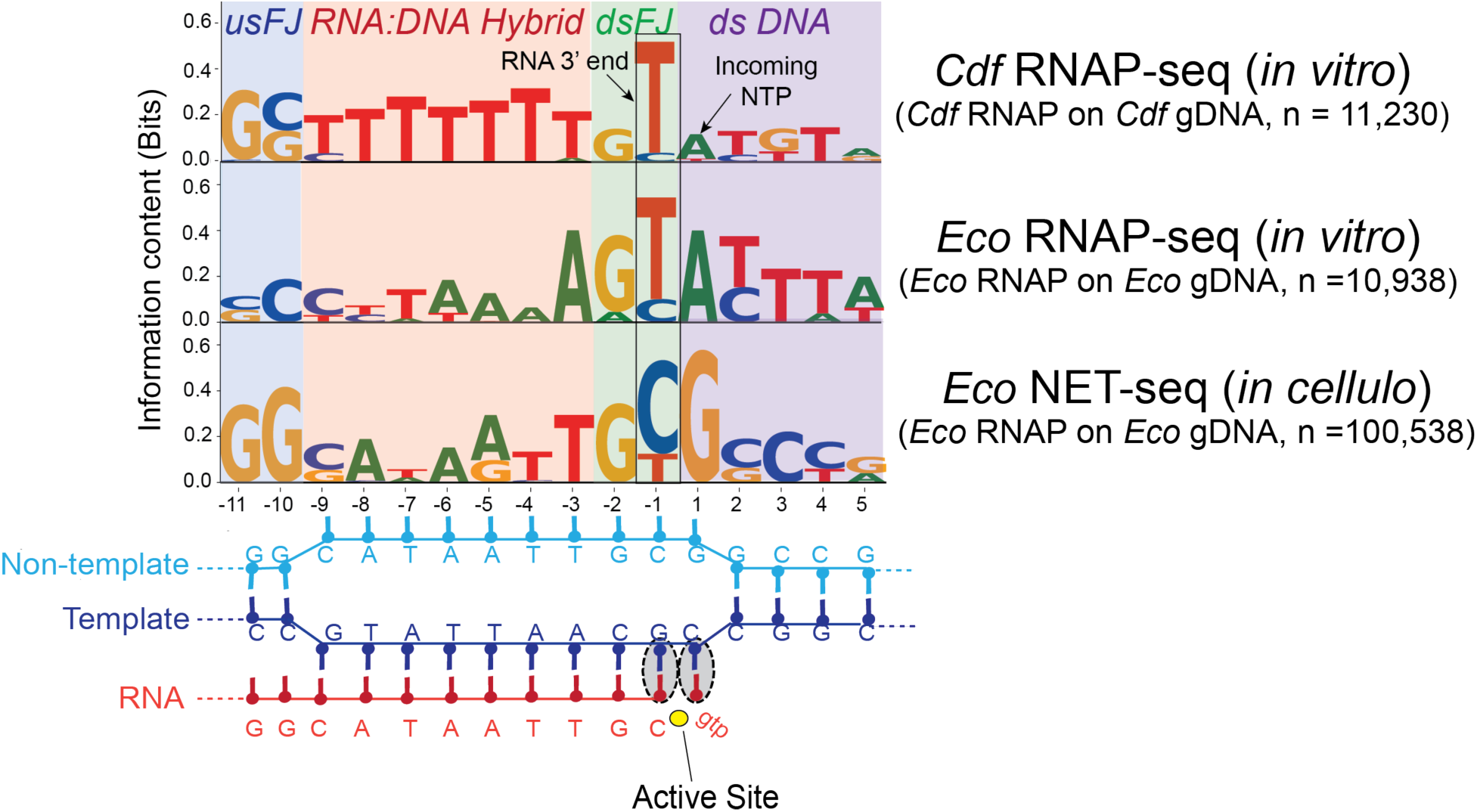
Consensus elemental pause sequences identified across RNAPs. Sequences were aligned at the RNA 3′ end (position -1) to generate consensus pause motifs spanning the transcription bubble (schematized below). Sequence logos were generated with genome-context normalization (Materials and Methods). Top, RNAP-seq using *Cdf* RNAP on *Cdf* gDNA; middle, RNAP-seq using *Eco* RNAP on *Eco* gDNA; bottom, *Eco* NET-seq (*in vivo*)^1^. The schematic below illustrates the transcription bubble, including upstream fork junction (usFJ), RNA:DNA hybrid, downstream fork junction (dsFJ), and downstream DNA (dsDNA) relative to the RNAP active site.

Although RNAP-seq does not distinguish between pausing and arrested states, these states likely represent a continuum of RNAP dwell times some of which are longer than the 15 min reaction time rather than reflecting completely distinct mechanistic classes. After excluding termination-associated sites, the remaining signals produce a consensus sequence highly similar to the ce-pause motif (Figure 3), suggesting that RNAP-seq predominantly captures bona fide pausing events. For clarity, we therefore refer to these non–terminator-associated halt sites as “pause sites” throughout the remainder of the study.

Together, these results demonstrate that elemental pausing is a conserved and intrinsic property of bacterial RNAPs across phylogenetically diverse species with markedly different genomic compositions, from the ∼50% A/T genome of *Eco* to the ∼70% A/T in *Cdf*.

### RNAP-seq reveals that *Cdf*NusG globally enhances transcriptional pausing and termination

To investigate the ability of RNAP-seq to characterize genome-wide TF activity, we next asked how *Cdf*NusG affected pausing by *Cdf*RNAP. NusG, the only universally conserved TF in all domains of life, remarkably exhibits divergent effects on pausing in different bacterial species^27^. In *Eco*, NusG suppresses transcriptional pausing^7,28^, whereas in *B. subtilis* and *M. tuberculosis*, NusG promotes pausing^9,10^. The genome-wide activity of *Cdf*NusG is unknown. Studying *Cdf*NusG function *in vivo* is difficult because it is essential and cannot be deleted^29,30^; knockdown approaches are confounded by its pleiotropic roles and functional coupling with NusA^31–34^. RNAP-seq provides a controlled *in vitro* system to directly assess NusG activity without these *in vivo* complications.

We performed RNAP-seq in the absence and presence of purified *Cdf*NusG and compared 3′-end halt scores genome-wide (including terminator-associated sites; Figure 4A). NusG increased a subset of halt signals across the genome (Figure 4B, Supplementary Table 5), indicating enhanced accumulation of RNAP at specific sites. We categorized halt sites into three groups: (1) sites unaffected by NusG; (2) sites shared between −NusG and +NusG conditions; and (3) sites uniquely detected in the presence of NusG (Figure S5A). Sequence logo analysis revealed no clearly distinct sequence features among these categories (Figures S5B–S5D).

**Figure 4:**
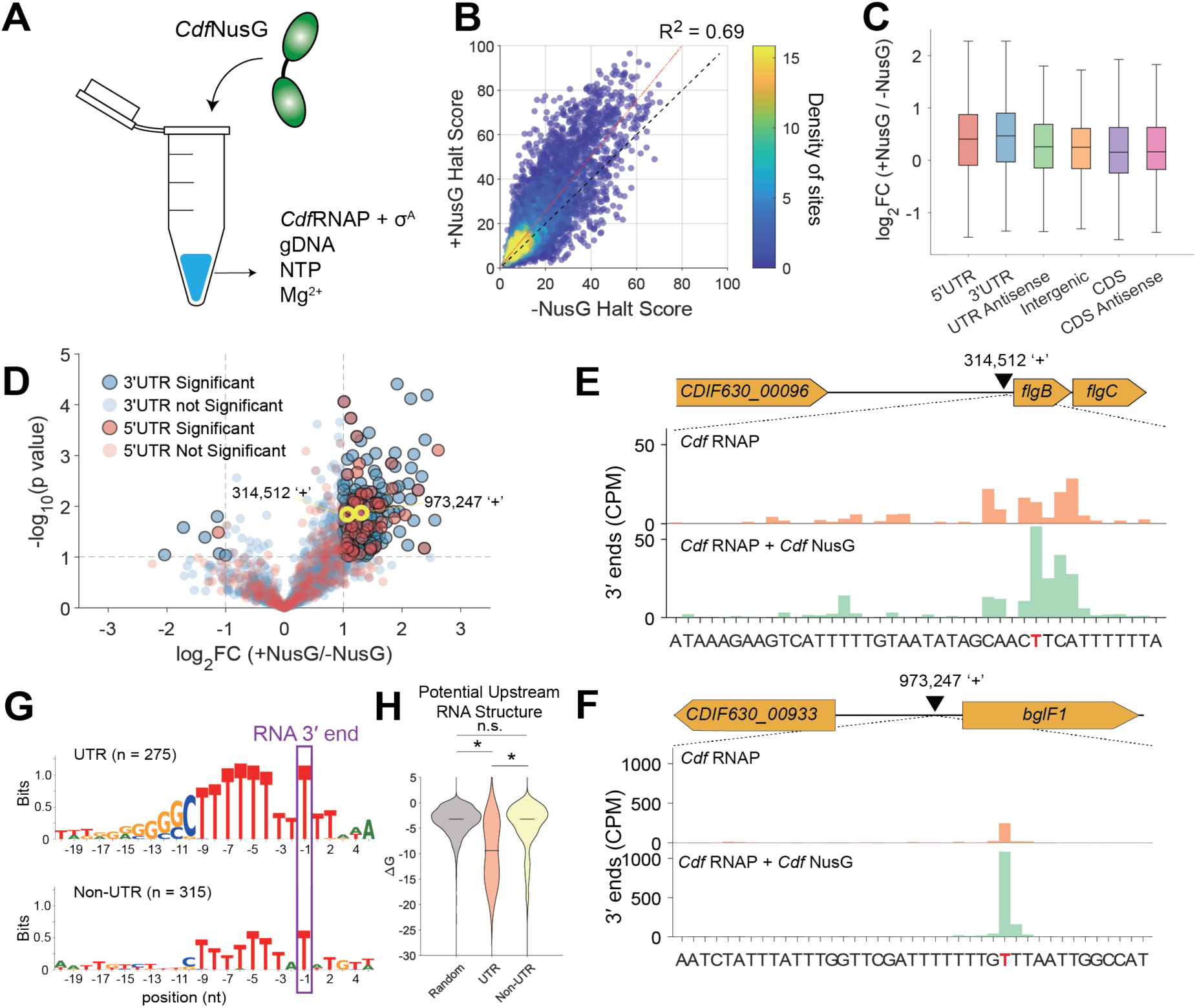
*Cdf*NusG enhances *Cdf*RNAP halting, with strongest effects at UTR regions. **A,** Schematic of RNAP-seq in the presence of NusG. **B**, Genome-wide comparison of 3′ end halt scores (n = 9494) of *Cdf*RNAP in the absence and presence of *Cdf*NusG. Each point represents a halt site; color indicates site density (heat map). The dashed line shows linear regression (R² = 0.69), revealing a global increase in halt scores upon NusG addition. Halt score is defined as the read count at each site normalized to the mean read count within a ±100 nt window. **C**, NusG-dependent changes in halt scores, shown as log₂ fold change (+NusG/-NusG). Box plots include 5′ UTRs (n=429), 3′ UTRs (n=1179), UTR antisense regions (n = 623), internal coding regions (n=1372), coding sequence (CDS; n=2821), and CDS antisense (n = 3168). The strongest enhancement occurs in 5′ and 3′ UTRs. **D,** Volcano plot of NusG-dependent changes in UTRs. Each point represents a halt site. The X-axis shows the log₂ fold change in halt score (+NusG/−NusG), and the y-axis shows statistical significance (−log₁₀ *p* value). Sites located in 5′ UTRs (red, n=429) and 3′ UTRs (blue, n=1179) are highlighted. Sites meeting significance thresholds (*p* < 0.1 and log₂ fold change > 1) are outlined in black (n=62 for 5′ UTR and n = 213 for 3′ UTR). Two representative sites (yellow circles) are shown in panels E and F. **E-F,** Representative loci illustrating NusG-enhanced halt signals in 5′ UTRs. The position of the RNAP halt site is indicated by a red nucleotide in the sequence. CPM-normalized signals are shown for *Cdf*RNAP alone (top) and in the presence of *Cdf*NusG (bottom), with gene annotations and sequence context. **G**, Sequence logos of halt sites in the presence of NusG, shown for UTR (top) and non-UTR (bottom) regions. Logos span −20 to +5 relative to the RNA 3′ end (-1). UTR-associated sites display a characteristic T-rich tract characteristic of intrinsic terminators, whereas non-UTR sites lack this feature. **H**, Predicted RNA secondary structures upstream of halt sites in the presence of NusG. Minimum free energy (ΔG) was calculated for sequences within 40 nt upstream of each RNA 3′ end using RNA-fold ^42^. UTR-associated halt sites exhibit significantly more stable predicted hairpin structures (more negative ΔG) compared with non-UTRs and random genomic sequences. (Mann–Whitney U test; n.s., *p* = 0.15;****, *p* < 0.0001)

Focusing on shared sites (detected in both +NusG and –NusG conditions, group 2), we found that NusG increases in 3′ end halt score were most pronounced in 5′ UTR and 3′ UTR regions (Figure 4C), with many sites showing >2-fold enhancement (Figures 4D–4F). Notably, NusG-enhanced sites in UTRs displayed a strong pattern of a predicted RNA hairpin followed by T-rich tracts, consistent with intrinsic terminator-like sequences (Figures 4G and 4H). Together, these results indicate that *Cdf*NusG acts as a global enhancer of RNAP halting, with particularly strong effects at UTR-associated regulatory regions.

Given that *Cdf* possesses unusually long 5′ UTRs (many >300 nt), in contrast to the shorter 20–60 nt 5′ UTRs typical of *Eco*^19^, we hypothesized that these regions may be enriched in regulatory RNA elements such as riboswitches. Previous studies have shown that NusG-dependent pausing can modulate riboswitch function co-transcriptionally^35^. These observations suggest the possibility that *Cdf*NusG also functions in these regulatory elements in 5′ UTRs. Consistent with this hypothesis, we identified 17 NusG-enhanced halt sites that overlap with previously annotated riboswitches^19^ (Figures S6A and S6B). Strikingly, we also detected 22 additional NusG-enhanced halt sites within 5′ UTR regions that lack prior annotation (Figure S6A, Supplementary Table 6). Rfam analysis of these loci revealed well-defined RNA secondary structures characteristic of regulatory RNAs^36,37^, suggesting that they represent previously unrecognized riboswitch-like elements.

Together, these findings indicate that *Cdf*NusG preferentially enhances RNAP halting within regulatory RNA regions and uncover a potential role for *Cdf*NusG in modulating both known and novel RNA-based regulatory elements in extended 5′ UTRs.

### *Cdf* RNAP preferentially pauses on antisense strands, in contrast to *Eco* RNAP

To investigate possible explanations for the difference in *Cdf*RNAP and *Eco*RNAP pausing behaviors (Figure 3), we performed RNAP-seq using both enzymes transcribing *Cdf* gDNA. We first quantified pausing fractions (defined as the proportion of total RNAP pause sites within each genomic feature) across the genome (Figures 5A, 5B). Overall pausing distributions were similar between the two RNAPs, with the notable exception of coding regions (CDS) and their corresponding antisense strands. Strikingly, *Cdf*RNAP exhibited a strong bias toward pausing on CDS antisense strands (− strand) (Figure 5A), whereas *Eco*RNAP preferentially paused on sense (coding, + strand) regions (Figure 5B). To determine whether this difference is reflective of underlying genome sequence features, we analyzed the nucleotide composition across the *Cdf* genome. We found that coding strands are enriched in purines (A+G), whereas antisense strands are enriched in pyrimidines (T+C) (Figure 5C), reflecting the overall AT-rich nature of the *Cdf* genome.

**Figure 5:**
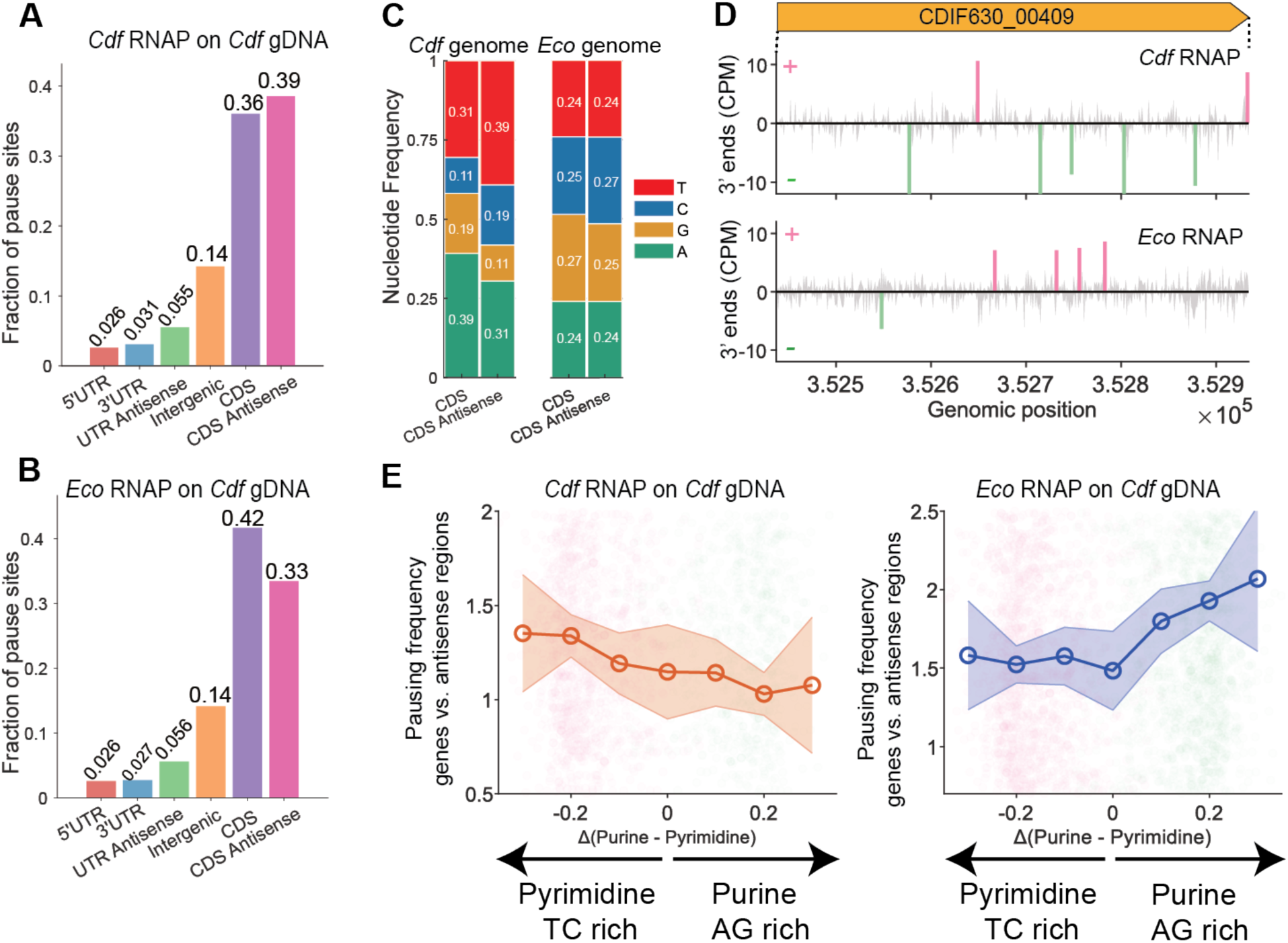
Genome-encoded purine–pyrimidine asymmetry drives RNAP-specific transcriptional pausing. **A–B,** Genome-wide distribution of transcriptional pausing sites identified for *Cdf* RNAP (**A**) and *Eco* RNAP (**B**) on the *Cdf* genome. Bar plots show the fraction of pause sites located in different genomic features, including 5′ UTRs, 3′ UTRs, antisense UTRs, intergenic regions, coding sequences (CDS), and CDS antisense regions. **C,** Nucleotide composition of CDS and their corresponding antisense regions on the *Cdf* and *Eco* genomes. Stacked bars indicate the relative frequencies of A, G, C, and T nucleotides within each region, with values indicated on the bars. **D**, Representative example of pausing sites within CDIF630_00409, located on the plus strand of the *Cdf* genome. Histograms show pause signals (counts per million, CPM) generated by *Cdf* RNAP (top) and *Eco* RNAP (bottom). Pause sites on the sense coding strand (+) are highlighted in red, whereas pause sites on the antisense strand (-) are highlighted in blue. **E**, Relationship between transcriptional pausing frequency and local purine–pyrimidine sequence bias at genome-scale. Each point represents an individual gene (sense strand, green) or its antisense region (pink). Circles indicate mean values within bins (bin width = 0.06), and shaded areas represent ±3× the standard error of the mean (SEM). Pausing frequency (normalized to number of pausing sites per kb) was plotted against Δ(Purine − Pyrimidine) for CDS regions and their corresponding antisense regions on the *Cdf* genome. Δ(Purine − Pyrimidine) is defined as the difference between the fraction of purines (A + G) and pyrimidines (C + T) within each region. Left panel, *Cdf* RNAP; right panel, *Eco* RNAP.

Consistent with these sequence biases, an examination of representative genomic loci showed that *Cdf*RNAP paused more frequently on the *Cdf* antisense strand, whereas *Eco*RNAP exhibits more frequent pausing on the *Cdf* sense strand (Figure 5D). To extend this analysis genome-wide, we quantified pausing frequency (number of pausing sites per kb per gene) as a function of local sequence composition. *Cdf*RNAP exhibited increased pausing sites in pyrimidine-rich regions that are enriched on *Cdf* antisense strands of genes, whereas *Eco*RNAP preferentially paused in purine-rich regions, which are enriched on *Cdf* coding (sense) strands (Figure 5E).

Together, these findings indicate that RNAP pausing is governed by both enzyme-specific properties and genome sequence context, with *Cdf* RNAP pausing more frequent at pyrimidine (TC)-rich sequences on antisense strand of genes, and *Eco* RNAP at purine (AG)-rich sequences on coding strands.

### Distinct homopolymeric sequence preferences underlie differential pausing by *Cdf* and *Eco* RNAP

To understand the sequence determinants of pausing by *Cdf*RNAP versus *Eco*RNAP more completely, we compared the pause sequence logos of the RNAPs transcribing either cognate or heterologous genomic templates. We focused our analysis on genes and their corresponding antisense regions. Specifically, we examined nucleotide composition within the −9 to −1 window relative to the RNA 3′ end (−1), corresponding to the transcription bubble and RNA–DNA hybrid. When transcribing the *Eco* genome, both RNAPs exhibited similar nucleotide compositions across pause sites, with approximately balanced A, T, G, and C frequencies (Figure 6A). In contrast, when transcribing the *Cdf* genome (*i.e.* genes and antisense genes), clear differences emerged: *Cdf*RNAP showed more enrichment of T residues within the −9 to −1 window, whereas *Eco*RNAP showed more enrichment of A residues in the same region (Figure 6B).

**Figure 6:**
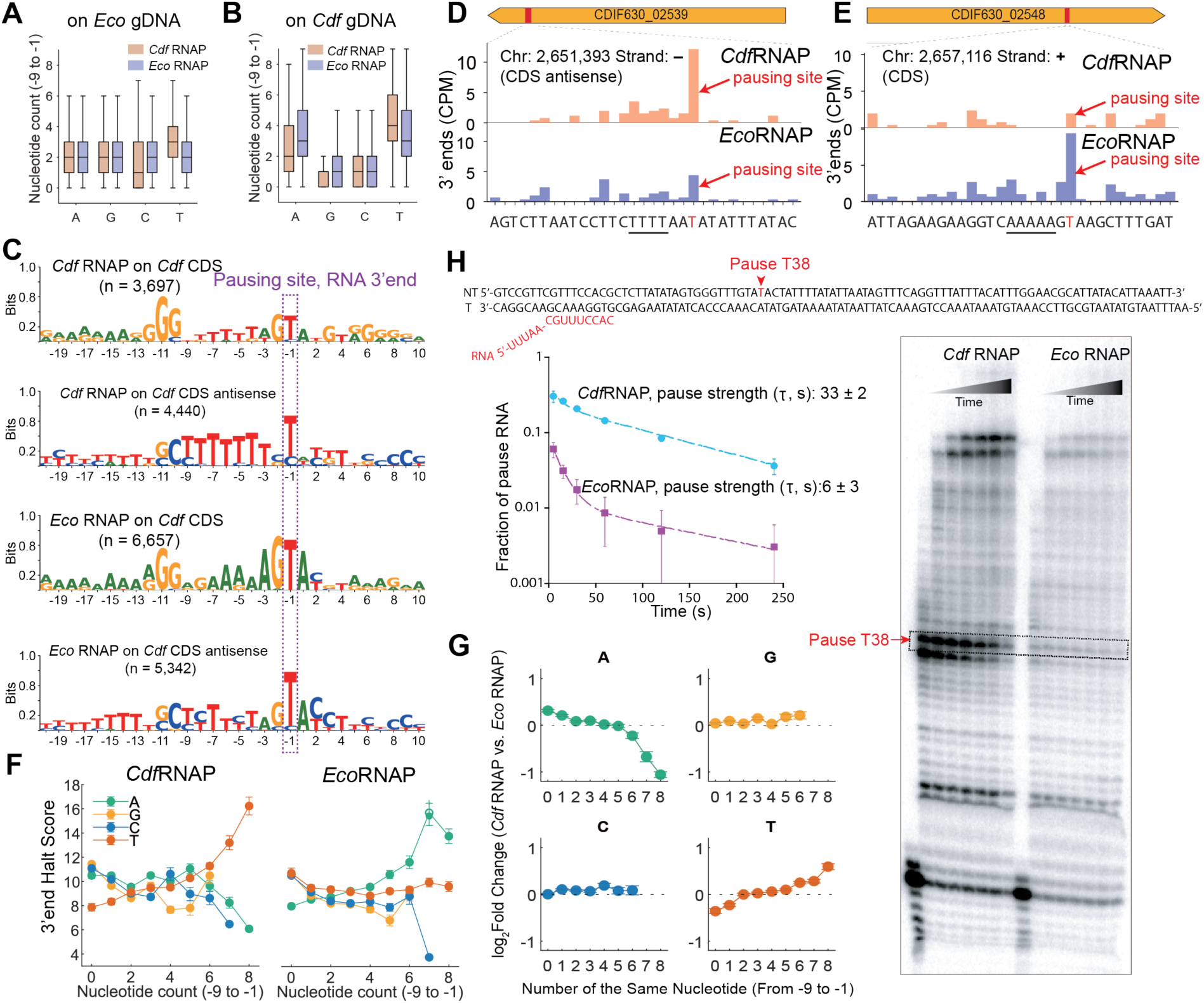
Nucleotide composition and homopolymeric enrichment in the −9 to −1 window influence differential RNAP pausing strength between *Eco* and *Cdf*. A-B, Nucleotide composition within the −9 to −1 nt window upstream of pause sites (intrinsic termination sites has not included). Boxplots show counts A, G, C, and T for pause sites identified by *Cdf*RNAP (orange) and *Eco*RNAP (blue). Panel **A** shows pause sites on the *Cdf* genome, panel **B** shows pause sites on the *Eco* genome. Boxes represent interquartile range (IQR) with median indicated; whiskers extending to 1.5×IQR. **C,** Sequence logos of pause sites generated by *Cdf*RNAP and *Eco*RNAP on both genomes. Logos depict nucleotide frequencies surrounding the pause position (-1), revealing distinct upstream sequence preferences depending on the RNAP and genomic context. **D-E**, Representative genes from the *Cdf* genome showing pause signals generated by *Cdf*RNAP and *Eco*RNAP on the same DNA templates. Histograms show CPM-normalized pause signals; paused positions are highlighted in red, and gene annotations are indicated above. **F**, Relationship between pausing strength and homopolymeric nucleotide enrichment upstream of pause sites. Mean pausing signal is plotted as a function of the number of identical nucleotides within the −9 to −1 window for *Cdf*RNAP (left) and *Eco*RNAP (right) on the *Cdf* genome. Error bars represent the standard error of the mean (SEM). **G**, Log₂ fold change in pausing strength (*Cdf*RNAP / *Eco*RNAP) as a function of the number of identical nucleotides in the −9 to −1 window. Increasing enrichment of specific nucleotides is associated with differential pausing behavior between the two RNAPs. H, *In vitro* transcription assays using a synthetic nucleic scaffold designed to test pausing activity of *Cdf* and *Eco*RNAP at a representative genomic site (*Cdf* genome, antisense strand −, position 2,414,338). RNA products from a time-course reaction were resolved by 12% denaturing urea–PAGE. “Chase” indicates reactions driven to completion by addition of high concentrations of NTPs (10 mM).

Consistent with this observation, sequence logo analysis of pause sites revealed distinct sequence preferences between the two RNAPs. *Cdf* RNAP preferentially pauses at T-rich sequences on antisense strands, whereas *Eco* RNAP preferentially pauses at A-rich sequences on sense strands (Figure 6C). Selected gene examples further illustrate these differences: *Cdf*RNAP exhibits stronger pausing at T-rich regions on antisense strands (Figure 6D), whereas *Eco*RNAP shows enhanced pausing at A-rich regions on coding (sense) strands (Figure 6E).

To quantify this relationship genome-wide, we analyzed pause strength (defined by the 3′-end halt score) as a function of homopolymer content within the −9 to −1 window. *Cdf*RNAP pause strength increased with the number of T residues, whereas *Eco*RNAP pause strength increased with the number of A residues (Figure 6F). In contrast, G and C content showed minimal effects on pausing for either enzyme.

We next directly compared pause strength between the two RNAPs by calculating the log₂-fold change (*Cdf*RNAP / *Eco*RNAP) as a function of nucleotide composition within the −9 to −1 window (Figure 6G). Consistent with the above trends, increasing T content strongly favored pausing by *Cdf* RNAP, whereas increasing A content favored pausing by *Eco*RNAP. No significant differences were observed for G or C content.

To validate the observed preference of *Cdf*RNAP for T-rich pause sites, we selected a representative T-rich site on the *Cdf* genome (antisense strand; position –2,414,338) and performed *in vitro* transcription assays using ³²P-labeled RNA as a readout (Figure 6H). The time course assay showed a discrete paused RNA species whose level decreased slowly over time after its appearance, which is hallmark of bona fide transcriptional pausing. Upon addition of high concentrations of NTPs (10 mM) in a chase step, nearly all the paused RNA elongated into longer transcripts, confirming that most RNAPs are paused and remain transcriptionally competent. A small fraction of the RNA persists after the chase, suggesting a minor population of arrested or termination-prone complexes form at the same site. This result is consistent with RNAP-seq result, which captures both pausing and more stable arrested states. Importantly, the dominant behavior at this site is pausing by *Cdf*RNAP, whereas *Eco*RNAP exhibits little pausing under the same conditions (Figure 6H).

Together, these results demonstrate that homopolymeric sequences within the transcription bubble differentially modulate pausing strength in a nucleotide- and RNAP-specific manner, with *Cdf*RNAP exhibiting stronger pausing at T-rich sequences, whereas *Eco*RNAP shows enhanced pausing at A-rich sequences.

## DISCUSSION

RNAP-seq provides a new platform to directly compare RNAP pausing, termination, and regulation of transcript elongation at genome-scale across species on both cognate and heterologous DNA templates in a controlled biochemical environment. This approach enables dissection of transcriptional pausing into its two primary determinants—DNA sequence context and trans-acting factors—without the confounding complexity of the cellular milieu. Using RNAP-seq, we identified distinct pausing signatures between *Eco* and *Cdf*RNAP and were able to distinguish contributions arising from intrinsic RNAP properties versus genome sequence context.

Both *Cdf*RNAP and *Eco*RNAP recognize versions of the previously defined ce-pause sequence^1,24–26,38^ in which the most conserved elements (GG or GC at –11,–10 and Y-R at –1,+1) are the same but in which other elements differ. Specifically, we found that *Cdf*RNAP pauses more frequently and more strongly at T-rich versions of the consensus pause sequence, whereas *Eco*RNAP preferentially pauses at A-rich versions of the consensus. Notably, these sequence preferences are preserved across both cognate and heterologous transcription conditions (Figure S7), indicating that they primarily reflect intrinsic properties of the RNAP enzymes rather than genome-specific effects.

It is currently unclear what differing features of *Cdf*RNAP and *Eco*RNAP account for this difference in pause-site recognition. Elemental pausing by *Eco*RNAP involves formation of multiple interconverting conformations of the elemental paused elongation complex^39^. In some of these conformations, the RNAP swivel domain rotates resulting in a blockage of the active site whereas others appear trapped in one of several pre-translocated states. It is possible that the energetics of these states differ in *Cdf*RNAP and *Eco*RNAP such that the pause sequence differences have different impacts on pausing overall. Examining the interactions of the two different consensus pause sequences with each RNAP by cryo-EM may yield insight into this question.

The *Cdf* genome exhibits pronounced strand asymmetry, with coding strands enriched in purines and antisense strands enriched in pyrimidines. Consistent with this organization, we observed that *Cdf*RNAP pauses more frequently and strongly on antisense strands. One potential biological implication is enhanced Rho-dependent termination on antisense transcripts^40^. Increased pausing may provide a temporal window for Rho to engage the nascent RNA and terminate transcription efficiently. Thus, the intrinsic pausing preference of *Cdf*RNAP may be functionally coupled to genome architecture to promote efficient antisense transcription termination (Figure 7). In contrast, *Cdf* coding strands are A-rich and exhibit reduced RNAP pausing. This reduced pausing may limit RNAP dwell time and thereby decrease opportunities for ribosome engagement, potentially contributing to the uncoupled nature of transcription and translation observed in some Firmicutes, such as *Bacillus subtilis* and possibly *C. difficile*^41^. However, the extent to which transcription–translation coupling is modulated by RNAP pausing dynamics in *Cdf* has not been tested directly as yet. More broadly, these results suggest that RNAP intrinsic properties and genome sequence composition have co-evolved to shape transcriptional regulation and gene expression programs in bacteria.

**Figure 7:**
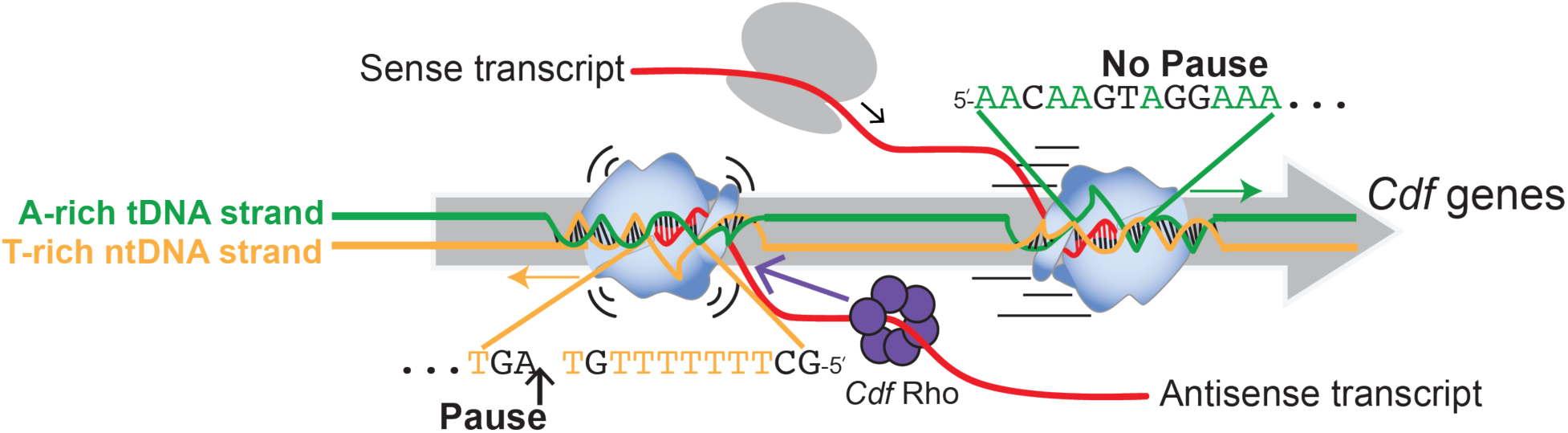
Schematic illustrating differential pausing of *Cdf*RNAP on coding (sense) and antisense strands of *Cdf* genes. Coding strands are enriched in A-rich sequences and exhibit reduced pausing, allowing efficient and continuous transcription elongation. This reduced pausing may potentially contribute to the uncoupled nature of transcription and translation in Firmicutes (*B. subtilis*, and possibly *C. difficile*)^41^. In contrast, antisense strands are enriched in T-rich sequences, which promote more frequent and longer-lived RNAP pausing. Consistent with previous findings (Dierksheide *et al*.)^40^, these extended pause states provide a temporal window for Rho to engage the nascent RNA and facilitate transcription termination on antisense transcripts.

RNAP-seq enables the discovery of regulatory features that are difficult to resolve *in vivo*. Using this approach, we identified *Cdf*NusG-enhanced pausing and termination sites within 5′ UTRs, including candidates for previously unannotated regulatory RNA elements, such as potentially novel riboswitches. These findings highlight the power of isolating transcription elongation factor activity at a genome-wide scale, which remains challenging in cellular contexts where multiple regulatory layers are intertwined. Future studies will be required to determine how *Cdf*NusG-mediated pausing and termination at 5’UTRs influence downstream gene expression and regulatory outcomes.

Notably, *Cdf*NusG functions as a clear pro-pausing factor, in contrast to NusG homologs in *E. coli*^7,8^. However, unlike *B. subtilis* NusG, which preferentially enhances pausing at TTnTTT motifs^9^, *Cdf*NusG does not exhibit this sequence specificity. This distinction suggests mechanistic divergence in NusG function across species and raises the possibility that *Cdf*NusG recognizes alternative sequence or structural features to modulate pausing.

Finally, RNAP-seq provides a versatile platform to compare transcriptional behavior across diverse RNAPs and their elongation factors. By enabling transcription on both cognate and heterologous genomic templates under defined conditions, it allows direct dissection of intrinsic RNAP properties and trans-acting regulatory mechanisms. This approach can be extended to study many more phylogenetically diverse RNAPs and transcription factors, offering a general framework to uncover conserved and lineage-specific principles of transcription regulation.

## ACKNOWLEDGEMENTS

We thank members of the Landick group for extensive discussions during the time the RNAP-seq method was developed at UW–Madison. Their invaluable insights and support made the results possible. We also thank Gene-Wei Li, Julia Dierksheide, and Robert Battaglia for helpful discussions about predicting intrinsic terminators and for sharing data results prior to publication. This work was funded by the National Institutes of Health (grant no. R00AI166036 to X.C. and R01GM038660 to R.L.), The Welch Foundation (grant no. I-2228 to X.C.) and The Endowed Scholars Program at the University of Texas Southwestern Medical Center (to X.C.). DNA sequencing services were provided by the UW–Madison Biotechnology Center DNA Sequencing Core Facility (RRID:SCR_017759).

## AUTHOR CONTRIBUTION

Conceptualization of experiments and analyses: PS, MW, JS, RL, XC; Methodology: PS, MW, RM, YB, EA; Investigation: PS, XC, RL; Writing: PS, XC; Editing: RL; Resources: RL, XC; Supervision: RL, XC.

## MATERIALS AND CORRESPONDENCE

Correspondence and requests for materials should be addressed to Robert Landick and Xinyun Cao.

## Supplementary Figures

**Figure S1.**
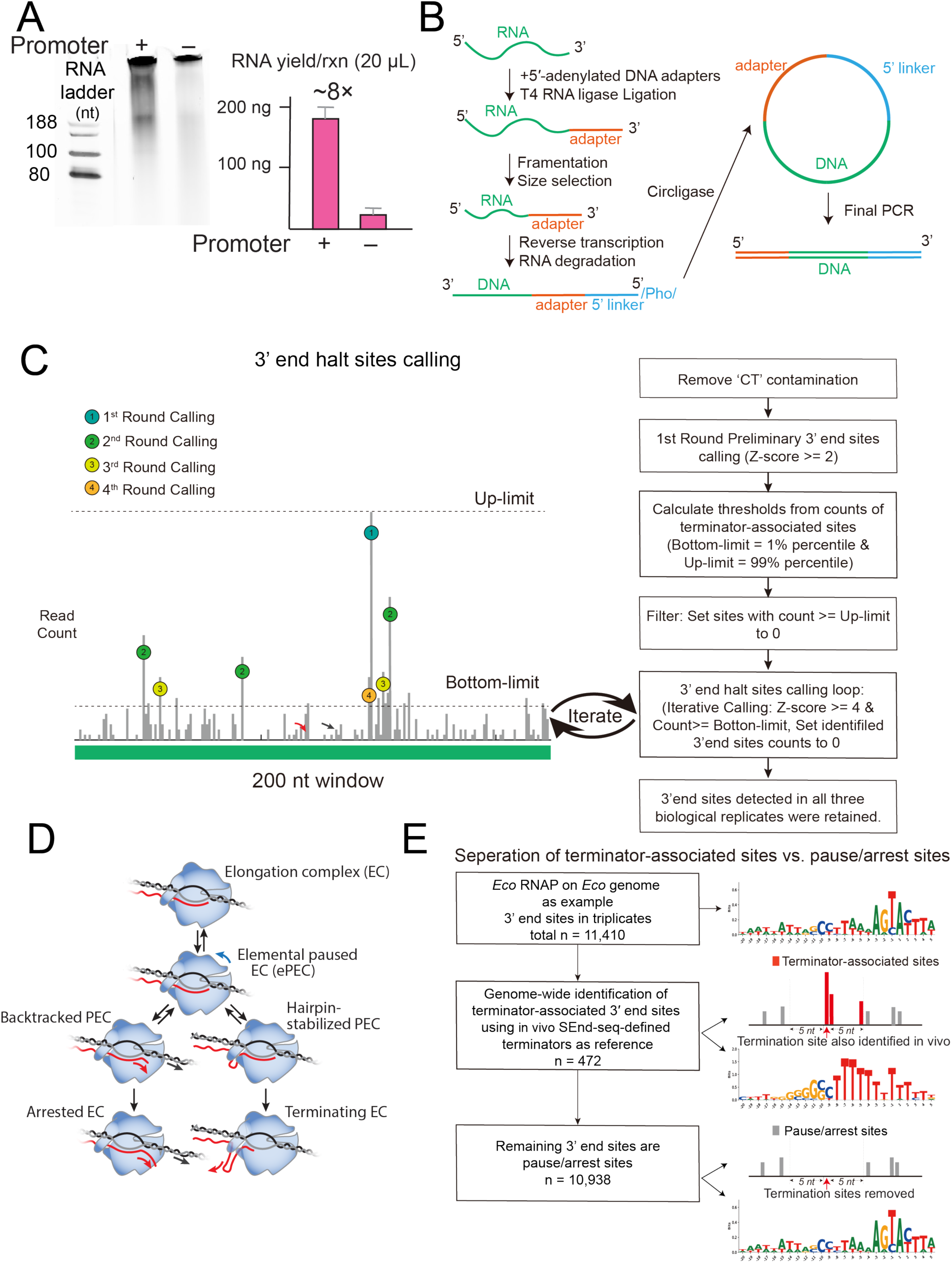
Pipeline for library preparation and transcript 3′ end halt site identification in RNAP-seq. **A,** Comparison of genome-wide transcription with and without promoter ligation using *Eco*RNAP transcribing *Eco* genomic DNA. Ligation of the strong *Cdf rrnC* promoter increases total transcript yield by ∼8-fold. **B,** RNAP-seq workflow for transcript 3′ end library preparation and sequencing. **C**, Pipeline to identify 3′ end halt sites generated by RNAP-seq. All samples were downsample to equilize sequencing depths (CPM). **D**, Illustration of kinetic transitions among pausing, termination, and halted elongation complexes. **E**, Bioinformatic pipeline for separating terminator-associated sites and pause/arrest sites, illustrated using *Eco*RNAP transcribing *Eco* gDNA.

**Figure S2:**
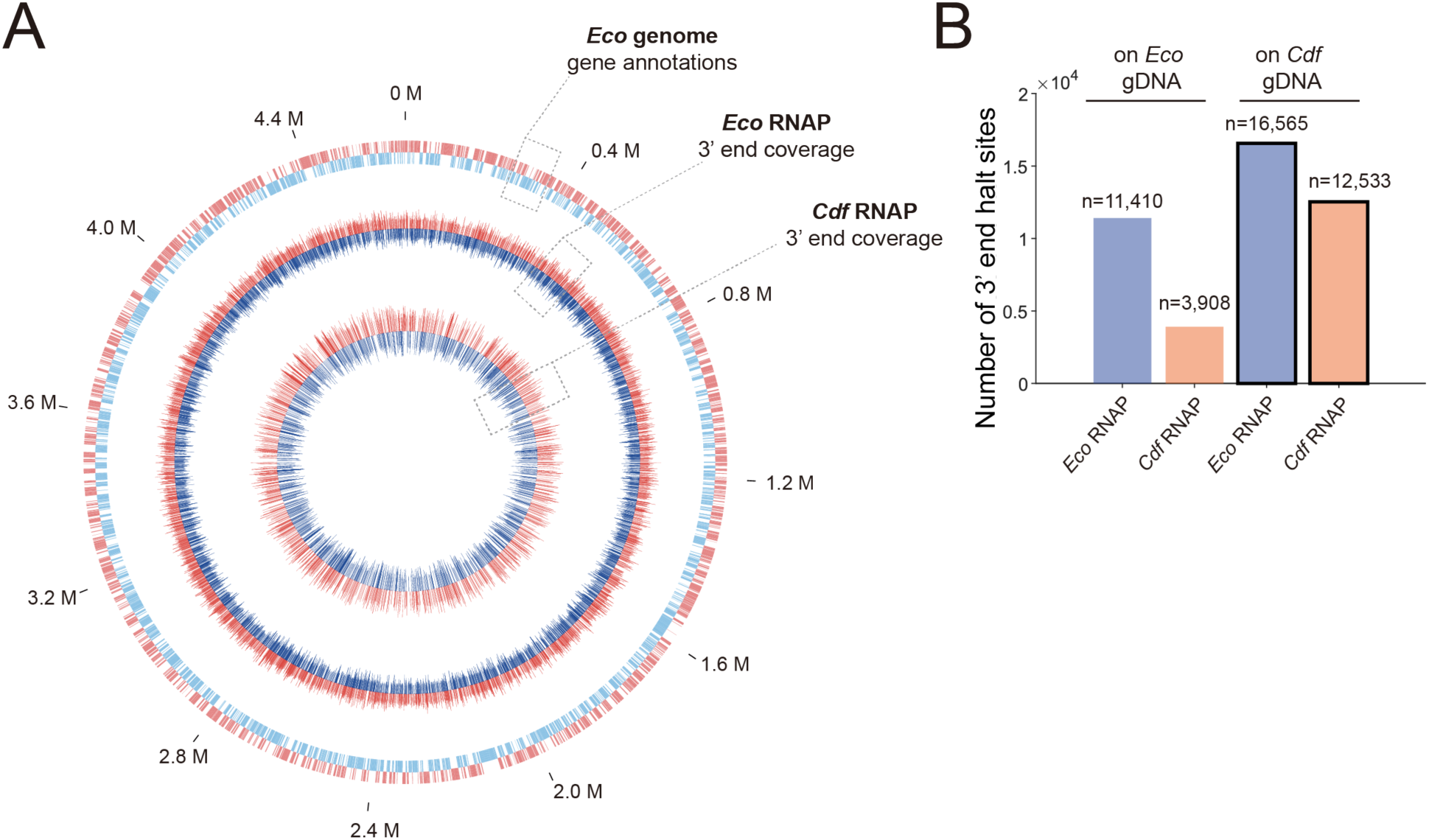
RNAP-seq identified transcript 3′ ends on the Eco genome. **A**. Genome wide distribution of 3′end halt signals on the *Eco* genome. Outer circle: gene annotations; middle circle: 3′ end coverage by *Eco* RNAP; inner circle: 3′ end coverage by *Cdf* RNAP. Red and blue colors represent positive and negative strands, respectively. Mb, megabase. **B**, Total number of halt sites detected across four conditions. Blue bars, *Eco* RNAP. Orange bars, *Cdf* RNAP.

**Figure S3.**
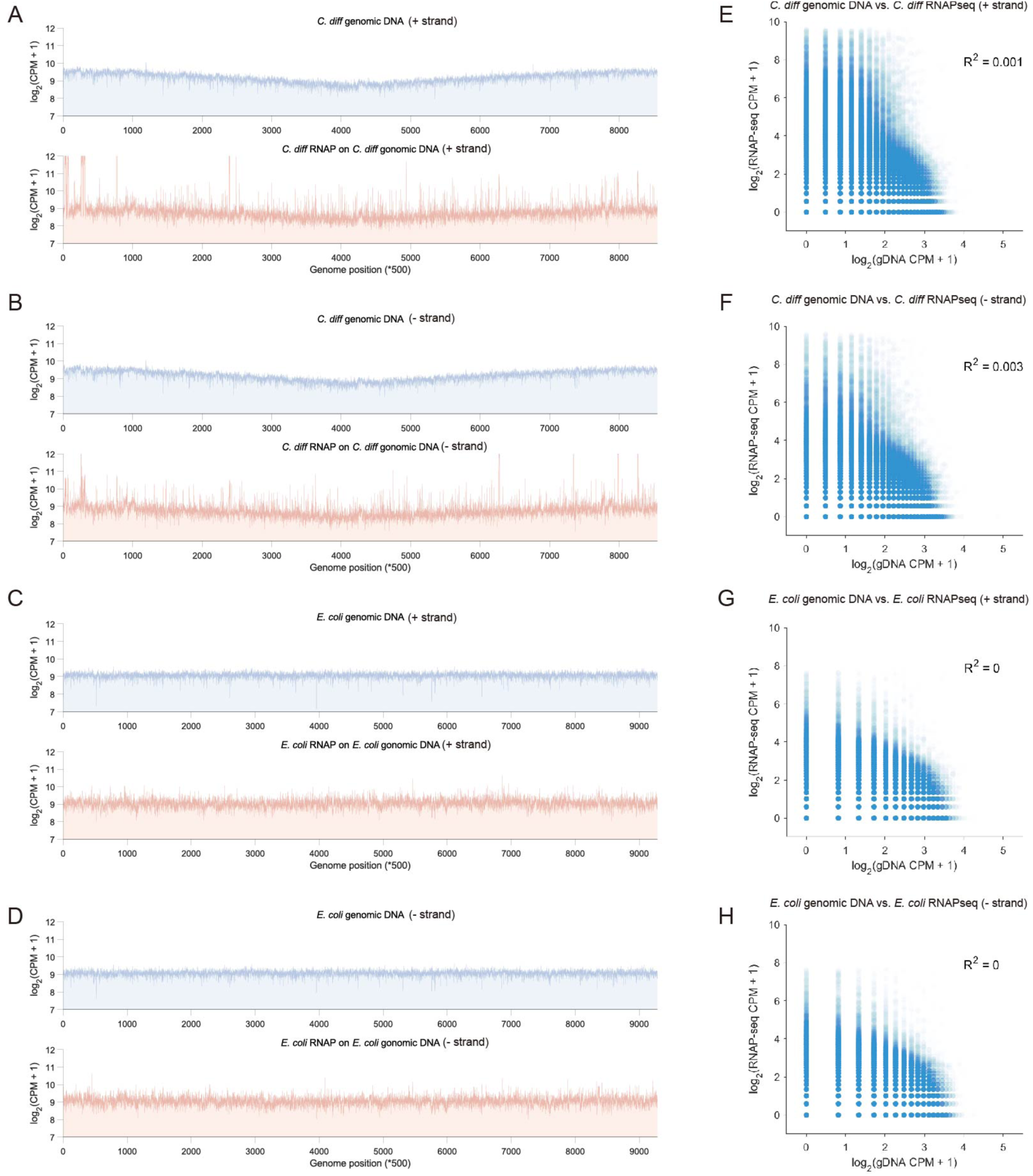
Genome-wide comparison of genomic DNA (gDNA) and transcript halt signals. **A–D**, Genome-wide distributions of log_2_-transformed gDNA 3′ end CPM and transcript 3′ end counts per million (CPM) across the *Cdf* or *Eco* genome, using 500-bp bins. For each bin, normalized counts were summed and plotted separately for the plus and minus strands. Panels A and B show gDNA and transcript halt signals for *Cdf* (plus and minus strands, respectively). The slight enrichment of gDNA signal near the origin reflects DNA isolated from actively growing cells. Panels C and D show gDNA and transcript halt signals for *E. coli* (plus and minus strands). gDNA 3′ end counts were derived from mapped gDNA fragment ends. Both gDNA and transcript signals were normalized to CPM prior to binning and log transformation. **E–H,** Scatter plots comparing log_2_ gDNA CPM and log_2_ RNAP-seq CPM at individual genomic positions for the corresponding datasets and strands shown in A–D. The very low coefficients of determination (R^2^) indicate that transcript signal is largely uncorrelated with the distribution of gDNA ends, demonstrating that transcript signal is not driven by DNA fragmentation or library preparation bias.

**Figure S4.**
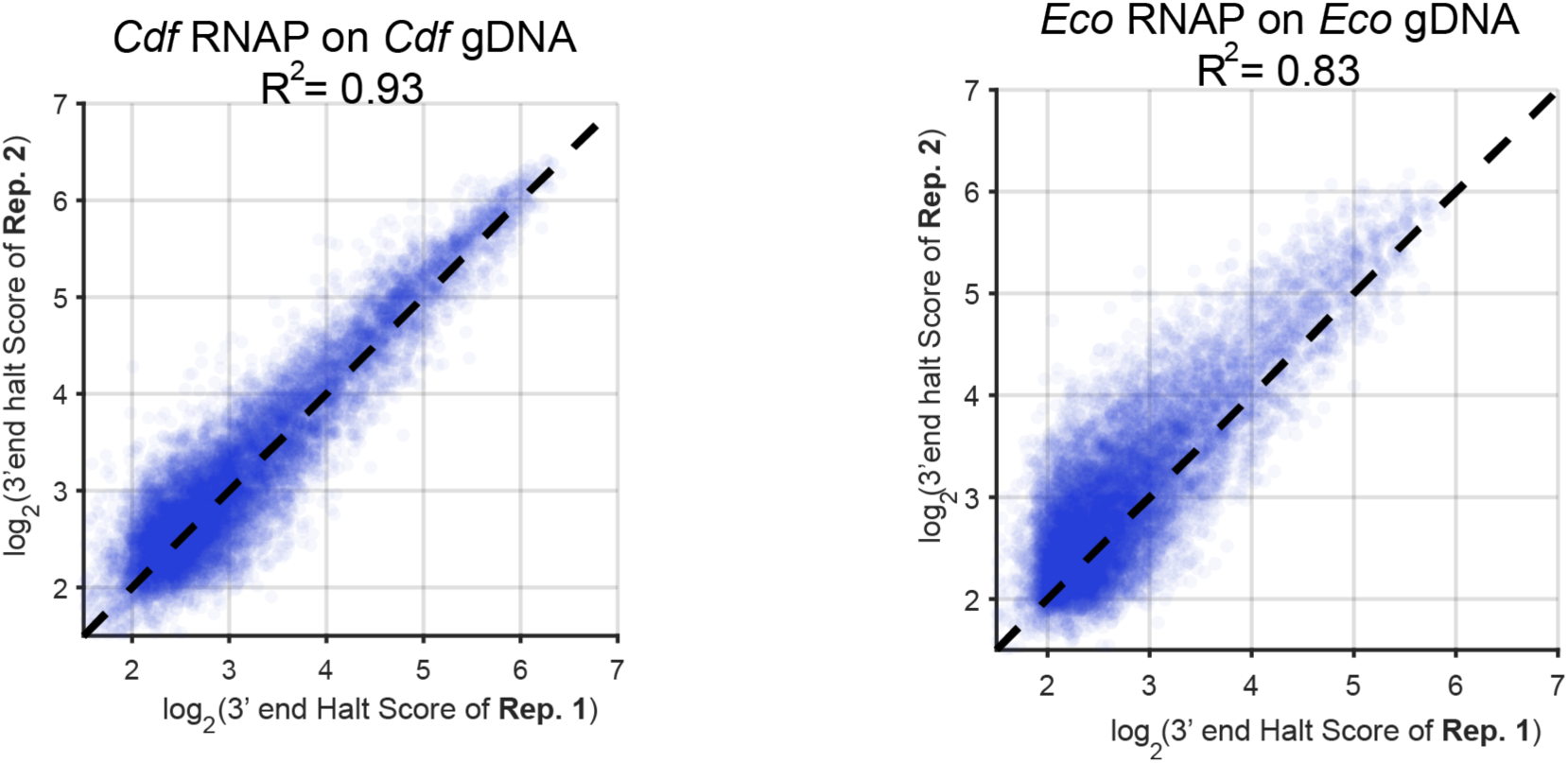
Genome-wide comparison of 3′ end halt score between two biological replicates. Reproducibility is assessed using Pearson’s correlation coefficient (R^2^). Each point represents an individual 3′ end halt site identified by RNAP-seq. The 3′ end halt score is defined as the read count at each halt site normalized to the mean read count within a 200-nt window centered on the site (±100 nt).

**Figure S5.**
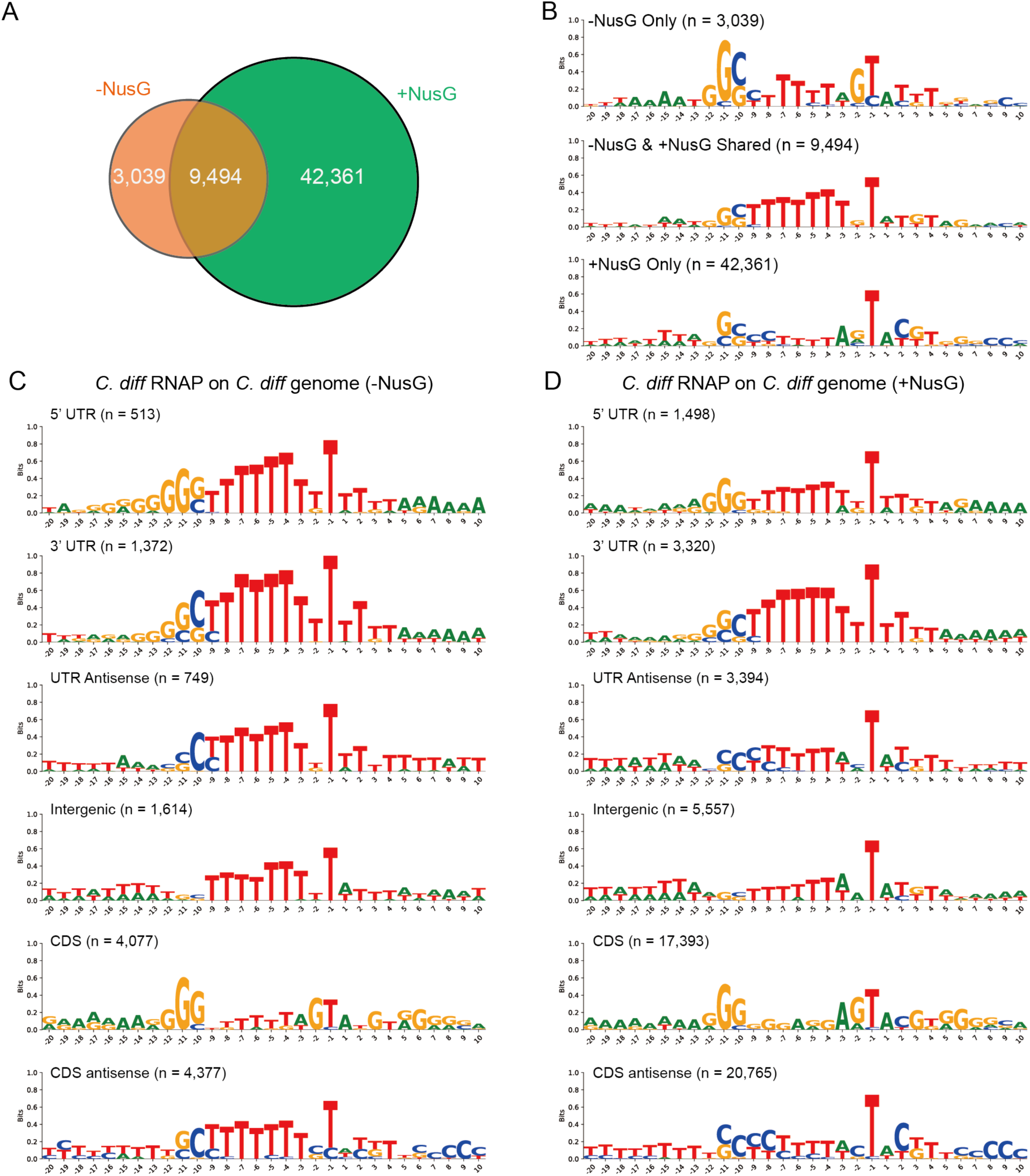
Sequence logos of NusG-enhanced halt sites on the *Cdf* genome **A,**Venn diagram showing the overlap of halt sites detected in the absence (−NusG) and presence (+NusG) of NusG. **B,** Sequence logos illustrating nucleotide enrichment surrounding 3′ end (−1) that are unique to −NusG, shared between conditions, or unique to +NusG. **C–D,** Sequence logos showing nucleotide features of halt sites across different genomic regions, including UTRs, antisense UTRs, intergenic regions, coding sequences (CDS), and antisense CDS. Numbers indicate the total number of sites in each category.

**Figure S6.**
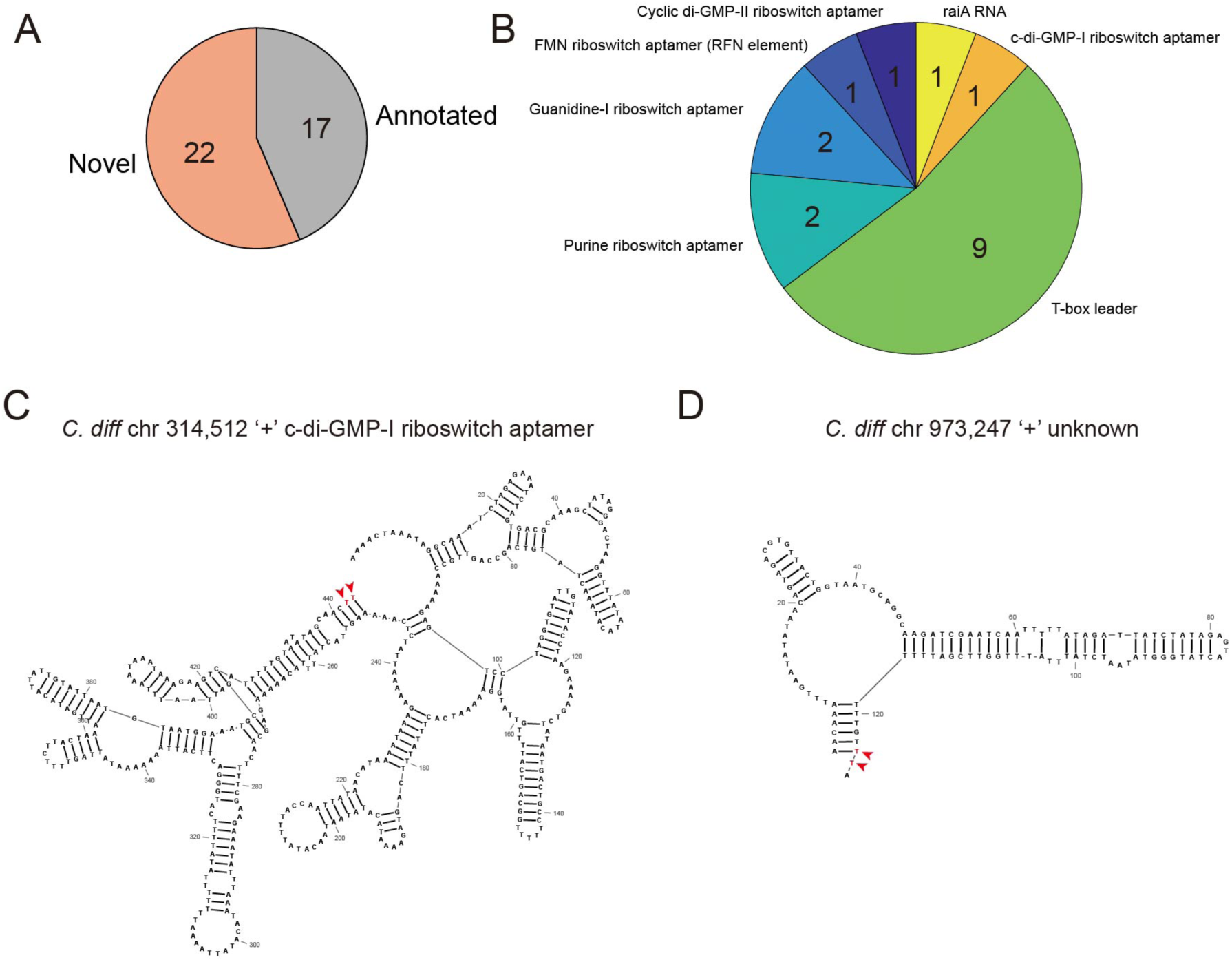
Analysis of NusG-enhanced halt signals in the 5′ UTR regions. **A,** Proportion of annotated versus novel halt sites enhanced by *Cdf*NusG. “Novel” sites lack annotation based on Rfam predictions^36,37^. For Rfam analysis, only transcripts with a length greater than 50 nt (from transcription start site to 3′ end) were included. **B,** Functional classification of *Cdf*NusG-enhanced halt sites with known regulatory RNA elements, including riboswitch aptamers and leader sequences. **C, D,** Putative RNA secondary structures in 5′ UTR regions of operons shown in main Figure 4E–F. RNA structures were predicted from the transcription start site to the RNA 3′ end. Arrows indicate the positions of halt sites.

**Figure S7.**
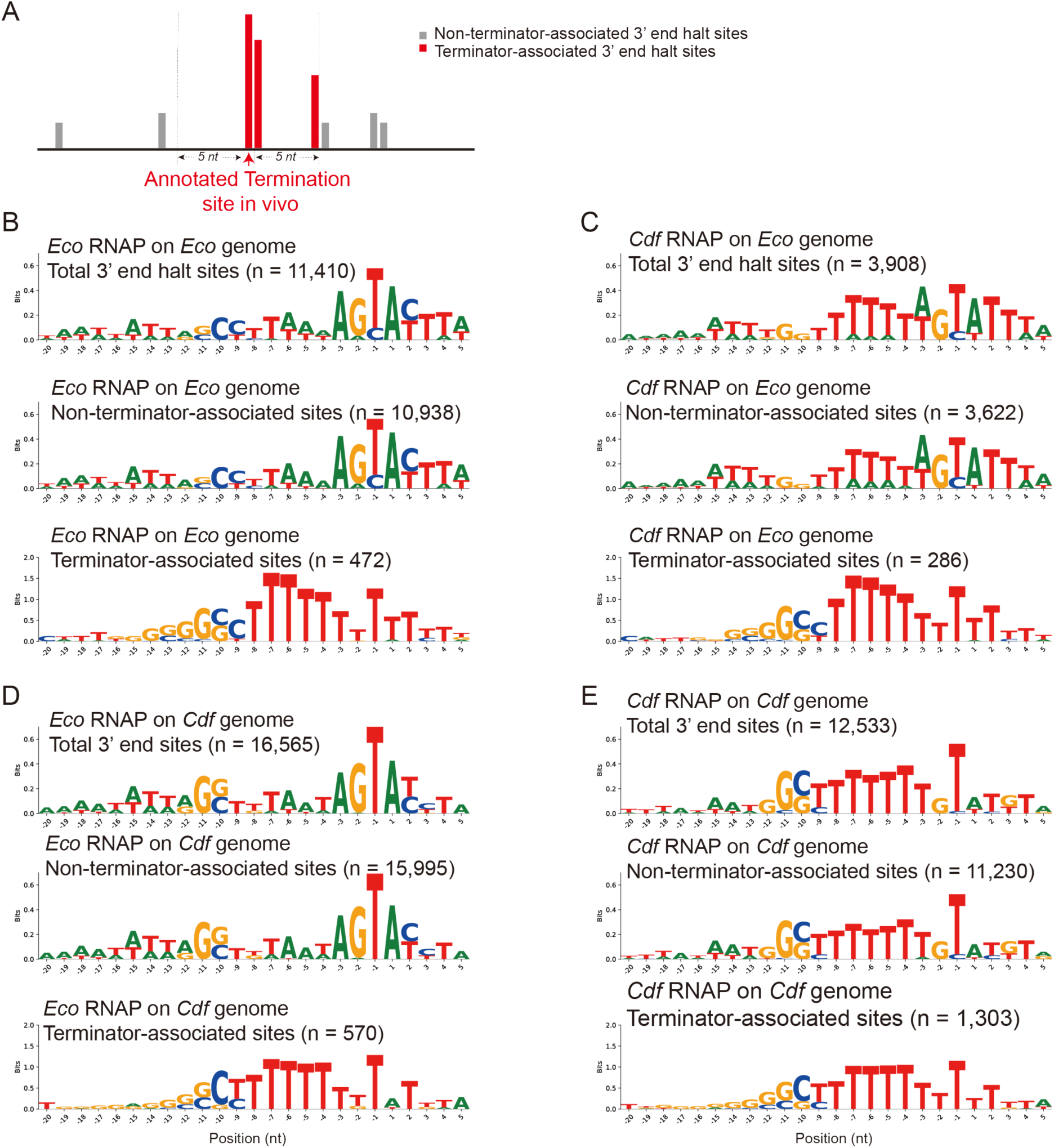
Sequence logos of terminator-associated and non–terminator-associated halt sites for *Eco* and *Cdf* RNAP transcribing cognate and heterologous genomic DNA

## SUPPLEMENTARY TABLES

Supplementary tables 1-6.

## MATERIALS AND METHODS

### Reagents

Antibiotics and chemicals used in this study were purchased from Sigma-Aldrich or Thermo-Fisher unless noted otherwise. [α-32P]GTP was obtained from PerkinElmer Life Sciences. Oligonucleotides were obtained from Integrated DNA Technologies.

### E. coli (Eco) cultures

*Eco* strains (MG1655 or BL21 DE3 derivatives) were grown in Luria–Bertani (LB) medium or on LB agar plates. When required, antibiotics or isopropyl β-D-1-thiogalactopyranoside (IPTG) were added.

### C. difficile (Cdf) cultures

*Cdf* strain CD630 was grown in an anaerobic chamber in brain heart infusion (BHI) medium (BD Difco; 37 g/L), supplemented with 5 g/L yeast extract and 0.1% L-cysteine, or on BHI agar plates containing the same supplements.

### Protein expression and purification *Cdf* α^A^

Cloning and purification of *Cdf* σ^A^ were performed as described previously^43^, with minor modifications. In brief, *Cdf* σ^A^ gene (KEGG: CD630_14550) from *Cdf* 630 was cloned into pET28a with an N-terminal His_10_ tag and expressed in *Eco* Rosetta^TM^ (DE3). Protein expression was induced with isopropyl-β-D-thiogalactopyranoside (IPTG) overnight at 16 °C. Cell pellets were resuspended in lysis buffer (50 mM Tris-HCl, pH 8.0, 1 mM EDTA, 5% glycerol, 5 mM DTT) and lysed by French press. Lysates were clarified by centrifugation (11,000 × g, 20 min, 4 °C, twice). The supernatant was subjected to Ni²⁺ affinity purification. The column was washed with 20 column volumes of wash buffer (20 mM Tris-HCl, pH 8.0, 0.5 M NaCl, 5% glycerol, 50 mM imidazole, and 1 mM DTT), and the His-tagged protein was eluted with 250 mM imidazole. Eluted protein was diluted 1.5-fold into TGEDZ buffer (10 mM Tris-HCl, pH 8.0, 5% glycerol, 0.1 mM EDTA, 5 µM ZnCl₂, 10 mM DTT) and further purified by heparin affinity chromatography (heparin FF 5 ml, Cytiva). After washing with 5mM NaCl TGEDZ buffer, σ^A^ was eluted using a linear NaCl gradient (up to 1 M NaCl) in TGEDZ buffer, and peak fractions were pooled and concentrated. The protein was stored in 20 mM Tris-HCl, pH 8, 10% (v/v) glycerol, 0.3 M NaCl, and 5 mM DTT at –80 °C.

### Eco σ^70^

*Eco* σ70 gene (*rpoD,* KEGG: b3067) from *Eco* K-12 MG1655 was cloned into pET28a with an N-terminal His_10_ tag and expressed in *Eco* BL21DE3. Protein expression and purification were performed as described above for *Cdf* σ^A^.

### *Cdf* NusG

Cloning and purification of *Cdf* NusG were performed as previously described with minor medications^44^. The *Cdf nusG* gene (CD630_00600) was PCR amplified from strain CD630 genomic DNA. The PCR project was then cloned into the pTYB2 vector (Addgene, catalog No. N6702S) using NEB HiFi DNA assembly to generate a C-terminal fusion with the Saccharomyces cerevisiae VMA intein and chitin-binding domain (CBD). The construct was transformed into *Eco* B834 (λDE3) (Novagen) cells for protein expression. Cells were grown in LB medium and protein expression was induced with 0.3mM IPTG at 16 °C for overnight (∼18 hours). Cell pellets were resuspended in chitin wash buffer (30 mM Tris-HCl, pH 8.0, 0.5 M NaCl, 1 mM EDTA, 0.05% Tween-20) plus one tablet of Roche cOmplete ULTRA EDTA-Free protease inhibitor. The cell suspension was lysed by sonication. Lysates were clarified by centrifugation (11,000 × g, 15–30 min, 4 °C). The supernatant was loaded onto a column (BioRad, Catalog No. 7321010EDU) packed with 3 ml of chitin resin pre-equilibrated with chitin wash buffer. The column was washed three times with 10 ml of chitin wash buffer, followed by on-column intein cleavage using 1.5 ml of cleavage buffer (chitin wash buffer supplemented with 75 mM DTT) at room temperature for overnight. Following cleavage, tag-free *Cdf* NusG protein was eluted using chitin wash buffer containing 10 mM DTT. Eluted fractions were pooled, injected into heparin affinity chromatography. After washing with buffer containing 10 mM Tris-HCl, pH 8.0, 5 mM NaCl, 5% glycerol, 0.1 mM EDTA, and 2 mM DTT, NusG was eluted using a linear NaCl gradient (up to 1 M NaCl) in the same buffer. Peak fractions were pooled and concentrated. The purified NusG was stored in 20 mM Tris-HCl, pH 8, 10% (v/v) glycerol, 0.3 M NaCl, and 5 mM DTT at –80 °C.

### *Cdf* RNA polymerase (RNAP)

Cloning and purification of *Cdf* RNAP were performed as described previously ^43^. Briefly, the *rpoA*, *rpoZ*, *rpoB*, and *rpoC* genes from *Cdf* 630 were codon-optimized for expression in *Eco*, assembled into a pET21-based vector, and engineered to encode a His-tagged RNAP complex. The β and β′ were fused using a polypeptide linker (LARHGGSGA) to ensure ensure 1:1 stoichiometry and inhibit assembly with the host *Eco* subunits. *Cdf* RNAP was overexpressed in *Eco* B834(λDE3) (Novagen) after induction with 0.3 mM IPTG in LB medium with 50 µg ml^−1^ kanamycin for overnight at 16 °C. Cells were lysed by French press in lysis buffer (50 mM Tris-HCl, pH 8.0, 1 mM EDTA, 5% glycerol, 5 mM DTT, 1× protease inhibitor cocktail (Halt Protease Inhibitor Cocktail (100 ×), Thermo Fisher), and 1 mM PMSF), and lysates were clarified by centrifugation (11,000 × g, 20 min, 4 °C, twice). RNAP were precipitated from the supernatant by gradual addition with mixing of polyethyleneimine (PEI) to 0.6% w/v final concentration, then washing three times with 10 mM Tris-HCl, pH 8, 0.25 M NaCl, 0.1 mM EDTA, 5 mM DTT, and 5% (v/v) glycerol , eluted with same buffer with 1M NaCl, and further purified by ammonium sulfate (final concentration to 35% w/v) precipitation. After centrifugation at 11,000g, 25 min, 4 °C, the RNAP was resuspended in 20 mM Tris-HCl, pH 8, 5% (v/v) glycerol, 0.5 M NaCl, and 5 mM β-mercaptoethanol and subjected to Ni^2+^ column purification. The column was washed with 20 column volumes of wash buffer (20 mM Tris-HCl, pH 8, 0.5 M NaCl, 5% (v/v) glycerol, 50 mM imidazole, and 5 mM β-mercaptoethanol) to remove contaminating proteins, and eluted with wash buffer containing 250 mM imidazole. The purified RNAP was concentrated, dialyzed into storage buffer (20 mM Tris-HCl, pH 8.0, 20% glycerol, 0.1 mM EDTA, 0.5 M NaCl, and 1 mM DTT), and stored at −80 °C.

### *Eco* RNAP

The *Eco* RNAP genes (*rpoA*, *rpoZ*, *rpoB*, and *rpoC*) from *Eco* K-12 MG1655 were cloned into a pET21-based vector engineered to encode a His_10_-tagged RNAP complex and expressed in *Eco* BL21(DE3). Protein expression and purification were performed as described above for *Cdf* RNAP.

### *Cdf* and *Eco* genomic DNA (gDNA) extractions and preparations for RNAP-seq

*Cdf and Eco* gDNA extractions were extracted according to a previously published protocol with minor modifications^45^. Briefly, *Cdf* was grown overnight in 10 ml BHIS medium at 37 °C under anaerobic conditions. *Eco* MG1655 (strain number RL3000) were grown overnight in 10 ml LB at 37 with vigourus shaking. Cells were harvested by centrifugation (4,000 × g, 10 min, 4 °C), washed once with 1ml TE (10 mM Tris-HCl pH8.0, 1 mM EDTA) buffer, and resuspended in 200 μl genomic DNA solution (34.23% sucrose (w/v) in TE buffer) supplemented with lysozyme (50 μl, 50 mg/ml). Samples were incubated at 37 °C for 1 h. Cell lysis was achieved by addition of 100 μl of 20% Sarkosyl and RNase A (15 μl, 10 mg/ml), followed by incubation at 37 °C for 30 min. Proteinase K (15 μl, 10 mg/ml) was then added and incubated for an additional 30 min at 37 °C. The volume was adjusted to 600 μl with TE buffer. DNA was extracted with an equal volume of phenol/chloroform/isoamyl alcohol (25:24:1, v/v/v), followed by centrifugation and transfer of the aqueous phase. This extraction step was repeated until no interphase was visible. The sample was extracted with chloroform to remove residual phenol. DNA was precipitated by 50 μl of 3 M sodium acetate (pH 5.2) and 3 volumes of cold 95% ethanol for overnight at -20. The next day pelleted by centrifugation at maximum speed 4 C. Finally, the DNA pellet washed with 500 μl cold 70% ethanol, and resuspended in 100 μl TE buffer.

For *in vitro* cell-free genomic assays, purified genomic DNA (gDNA) was sheared using a Covaris S220 Focused-ultrasonicator (ME220). Up to 15 μg of gDNA was loaded into a microTUBE-500 AFA Fiber Screw-Cap (Covaris) in 300 μl Tris–EDTA buffer (pH 8.0). DNA was sheared to an average fragment size of 0.5–1.5 kb using the following settings: duration, 80 s; peak power, 75; duty factor, 5%; cycles per burst, 100; and average power, 3.8. DNA nicks were repaired using the NEBNext® Ultra™ II End Repair/dA-Tailing Module (NEB, Cat. No. E7546L).

Fragmented *Eco* and *Cdf* gDNA (1 μg in TE buffer, adjusted to 50 μl) was subjected to end repair and 3′ dA-tailing using NEBNext® Ultra™ II End Repair/dA-Tailing Enzyme Mix (3 μl) and Reaction Buffer (7 μl) in a total reaction volume of 60 μl. The reaction was incubated at 20 °C for 30 min, followed by 65 °C for 30 min. The resulting dA-tailed gDNA was immediately subjected to promoter ligation (see next section).

### Promoter annealing and ligation to fragmented gDNA

Promoter oligonucleotides encoding the *Cdf rrnC* promoter were synthesized by IDT (Forward: 5′Biotin/TTTGACAAAAAAATATTTAAAATAAAAGTTAAAAAGTTGTTGACTTAGA ATAATATAGATGATATTATATATGAGTGAAAAGTAAATAGATAAATTAACT-3′; Reverse: 5′Phos/GTTAATTTATCTATTTACTTTTCACTCATATATAATATCATCTATATTATTC TAAGTCAACAACTTTTTAACTTTTATTTTAAATATTTTTTTGTCAAA-3′).

The forward and reverse oligonucleotides are fully complementary, except for a single 3′dT overhang on the forward strand to facilitate base pairing with dA-tail (see below). The −10 and −35 promoter elements are underlined in the sequences. Forward and reverse oligonucleotides (5 μM each) were then annealed in a 20 μl reaction containing 20 mM Tris-acetate (pH 8.0), 40 mM potassium acetate, and 5 mM Mg(OAc)₂. The mixture was heated to 95 °C and gradually cooled to 25 °C in a thermocycler to allow duplex formation.

Promoter ligation was performed by directly adding NEBNext Ultra II Ligation Master Mix (15 μl, Cat. No. E7546L), Ligation Enhancer (1 μl), and annealed promoter (1.5 μl, 5 μM) to the end-prepped DNA, followed by incubation at 20 °C for 15 min. Ligation products were purified using AMPure XP beads (0.7× followed by 0.9× size selection), washed with 80% ethanol, and eluted in nuclease-free water to remove excess promoter duplex. Successful promoter ligation was further verified using streptavidin magnetic beads (Thermo Fisher Scientific, Cat. No. 88817). Final promoter−gDNA concentration was quantified using the Qubit™ dsDNA Quantitation Broad Range assay (Thermo Fisher Scientific).

### Transcription reactions for RNAP-seq

*Cdf* (or *Eco*) RNAP holoenzyme was assembled by incubating core RNAP with σ^A^ or σ^70^ in transcription buffer (10 mM HEPES, 0.1 mM EDTA, 50 mM potassium glutamate, 10 mM Magnesium acetate, and 1 mM DTT) at 37 °C for 15 min. *In vitro* transcription reactions were carried out in a total volume of 200 µl containing assembled holoenzyme (0.375 µM), promoter−gDNA (25 nM), NTPs (1 mM each), transcription buffer, and RNase inhibitor (120 U, Promega, Cat. No. N2111). Reactions were incubated at 37 °C for 15 min. Transcription was stopped by degrading the DNA template with DNase (15 U, Promega, Cat. No. M6101) for 10 minutes at 37 °C. DNase was subsequently inactivated by adding an equal volume (215 μl) of phenol:chloroform:isoamyl alcohol (25:24:1, v/v/v), followed by phase separation. RNA was recovered by ethanol precipitation, resuspended in nuclease-free water, flash-frozen in liquid nitrogen, and stored at −80 °C until library preparation.

### Library preparations for capturing transcript 3′end

RNA libraries were prepared largely as described previously^12,20^, with minor changes. 5′-adenylated DNA adapters (5′-/Phos/CTGTAGGCACCATCAAT/3′-ddC, 6 μM) were generated using 6 μM Mth RNA ligase (NEB, Cat E2610L) in the presence of 80 µM ATP and 25% PEG 8000. Reactions were incubated at 65 °C for 4 h, followed by heat inactivation at 85 °C for 5 min. Purified RNA (300 ng–1 µg) from transcription reactions was ligated to 2 µM adenylated adapter using 6.7 U/µL T4 RNA ligase 2 (truncated; NEB, Cat. No. M0242L) in 1× T4 RNA Ligase Reaction Buffer supplemented with 25% PEG 8000 and 1.5 U/µL RNase inhibitor (Promega, Cat. No. N2111) at 37 °C for 3 h. Ligation reactions were terminated by addition of EDTA to a final concentration of 16 mM. Ligated RNA was fragmented by addition of an equal volume of alkaline buffer (12 mM Na₂CO₃, 88 mM NaHCO₃, 2 mM EDTA) and incubation at 95 °C for 35 min. RNA was purified by isopropanol precipitation in the presence of 0.3 M sodium acetate and GlycoBlue. Size-selected RNA fragments were resolved on a 15% TBE–urea polyacrylamide gel and recovered by gel extraction. Reverse transcription was performed using SuperScript III Reverse Transcriptase (Invitrogen, Cat. No. 18080044) in the presence of dNTPs and the RT primer: (5′/Phos/AGATCGGAAGAGCACACGTCTGAAC/iSp18/CACTCA/iSp18/CCTACACGACG C TCTTCCGATCTTCCGACGATCATTGATGGTGCCTACAG-3′) at 50 °C for 1 h. RNA was subsequently degraded by alkaline hydrolysis using 0.1 M NaOH at 98 °C for 20 min and neutralized with 0.1 M HCl. cDNA products were resolved and purified from a denaturing 10% TBE–urea polyacrylamide gel. Purified cDNA was circularized using CircLigase (Lucigen, Cat. No. CL9021K) at 5 U/µL in the presence of 50 µM ATP and 2.5 mM MnCl₂ at 60 °C for 1 h, followed by heat inactivation at 80 °C for 10 min. Circularized cDNA was amplified by PCR using Q5 DNA polymerase (NEB) with Illumina indexed primers, and products were size-selected on an 8% TBE polyacrylamide gel. Final library concentration and fragment size distribution were assessed using a High Sensitivity D1000 ScreenTape assay on the TapeStation system (Agilent). Libraries were sequenced at the University of Wisconsin–Madison Biotechnology Center on an Illumina NovaSeq X platform using 150-nt paired-end reads.

### 32P-labeled *in vitro* transcription assays

Elongation complexes (ECs) were assembled using a direct reconstitution approach. Pre-assembled nucleic-acid scaffolds (10 μM RNA and 20 μM template DNA (t-DNA)) were incubated with *Eco* or *Cdf* RNAP core enzyme in transcription buffer (20 mM Tris-acetate, pH 8.0, 40 mM potassium acetate, 5 mM Mg(OAc)₂, and 1 mM DTT) to yield final concentrations of 0.5 μM RNA, 1.0 μM t-DNA, and 7.5 μM RNAP. Complexes were assembled at 37 °C for 5 min, followed by addition of non-template DNA (nt-DNA) to 2.5 μM and further incubation for 5 min. Then 10mg/ml Heparin were added to 0.1 mg/ml and further incubation for 3 min., yielding ∼500 nM elongation complexes (ECs). For radiolabeling, ECs were incubated with 10 μCi [α-³²P]GTP (3000 Ci mmol⁻¹) for 5 min at 37 °C. Additional GTP was added such that the final concentration of GTP in the solution was 2.5 µM, and incubation continued for 3 min at 37 °C. Elongation was restared by addition of 2× NTPs (200 μM each). Time points were taken by mixing 5 µl reaction aliquots with 5 µl of 2× stop buffer (8 M urea, 50 mM EDTA, 1×TBE, pH 8.3, 0.02% bromophenol blue, and 0.02% xylene cyanol). “Chase” represents reactions driven to completion by addition of 10 mM NTPs. RNA products were resolved by 12% Urea-PAGE with 0.5× TBE running buffer and scanned by a Typhoon Phosphorimager. To quantify effects in ImageQuant, boxes were drawn at the pause band, before the pause band, and after the pause band. After background subtraction, the fractions of RNA at pause position was calculated, and values were averaged across three technical replicates; error bars reflect standard deviation. Pause kinetics were analyzed by fitting the time-dependent change in pause fraction to a bi-exponential decay model: y=F_fast_ × e ⁻^Kfast^ × ^t^ + F_slow_ × e ⁻^Kslow^ × ^t^, where *F*_fast_ and *F*_slow_ represent the fraction of RNA species entering the slow and fast escape phase and *k*_fast_ and *k*_slow_ are the corresponding escape rate constants. The pause strength (τ) was calculated as: τ= F_fast_/k_fast_ + F_slow_/k_slow_

### Fragmented genomic DNA sequencing for *Cdf* and *Eco*

Fragmented genomic DNA libraries were prepared using the NEBNext Ultra II DNA Library Prep Kit, fragmented genomic DNA was subjected to end repair and dA-tailing using the NEBNext Ultra II End Prep Enzyme Mix, followed by adaptor ligation with NEBNext adaptors. Ligated DNA was purified using AMPure XP beads and amplified by PCR using NEB Q5 polymerase and illumina indexed primers. Amplified libraries were size-selected, purified, and quantified prior to sequencing.

### Illumina sequencing & Data processing RNAP-seq data processing

Adaptor sequences were removed from raw FASTQ reads using Cutadapt^46^. Processed reads were aligned to the *Eco* MG1655 reference genome (NCBI Reference Sequence: NC_000913.3) or the *Cdf* 630 reference genome (NCBI Reference Sequence: CP010905.2) using Bowtie2^47^. Alignment files in SAM format were converted to BAM and BED formats using SAMtools^48^ and BEDtools^49^, respectively.

### Fragmented genomic DNA data processing

Genomic DNA reads were processed using the same pipeline as RNAP-seq data, except that adaptor trimming was not performed.

### 3′ end halt sites calling from RNAP-seq data

3′ end halt sites were identified from genome-wide RNAP-seq 3′ end profiles using an iterative peak-calling pipeline. Briefly, reads corresponding to adaptor-derived artifacts (‘CT’) were first removed, and 3′ end counts were obtained at single-nucleotide resolution. An initial set of candidate 3′ end peaks was defined using a Z-score threshold (Z ≥ 2). Peaks associated with intrinsic terminators were then identified by mapping to annotated terminator regions. Based on the distribution of 3′ end signals at terminator-associated sites, lower (1st percentile) and upper (99th percentile) count thresholds were determined. To reduce the influence of extreme outliers, sites with counts above the upper threshold were reset to zero. Refined 3′ end peaks were subsequently identified using more stringent criteria (Z ≥ 4 and count ≥ lower threshold). After each round (4 rounds total) of peak calling, identified peaks were removed from the signal track by setting their counts to zero, and the peak-calling procedure was repeated iteratively for four rounds to capture peaks across a broad dynamic range. Finally, only 3′ end peaks consistently detected across all biological replicates were retained as high-confidence sites for downstream analyses.

### RNAP-seq data analysis

#### Calculation of halting score for halt sites

For each genomic position 𝑖, a local halting score was defined as the ratio of read counts at position 𝑖 to the mean read counts in a local flanking window:

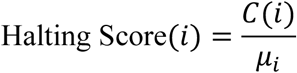

where 𝐶(𝑖)denotes the normalized read counts at position 𝑖, and 𝜇_1_ is the local mean read counts calculated within a ±100 nt window centered at position 𝑖.

### Sequence Logo with genomic background correction

Aligned sequences were used to compute position-specific nucleotide frequencies. To quantify positional enrichment relative to genomic background, information content was calculated using *Kullback–Leibler* (*KL*) divergence. Genome-wide background nucleotide frequencies were estimated separately for *Eco* and *Cdf* and used as reference distributions. For each nucleotide at each position, the information content (in bits) was calculated as

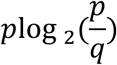

where *p* is the observed positional frequency and *q* is the corresponding genomic background frequency. Negative values were set to zero. Sequence logos were generated from the information matrices by using Python package *logomaker*.

### *Cdf* & *Eco* genome annotation

The *Cdf* 630 genome sequence (CP010905.2) and corresponding GFF3 annotations were used to define gene coordinates. Transcription start sites (TSSs) and transcription termination sites (TTSs) were obtained from published RNA-seq datasets^19^. Only high-confidence TSSs annotated as primary or secondary were retained. For each TSS, the nearest coding sequence (CDS) on the same strand was identified, and the 5′ UTR was defined as the region between the TSS and the CDS boundary. Identification of 3′ UTRs was performed analogously by assigning each TTS to the nearest upstream CDS on the same strand and defining the intervening region as the 3′ UTR. UTRs longer than 500 nt were excluded. Antisense UTRs were defined as the genomic regions antisense to annotated 5′ and 3′ UTRs. Intergenic regions were defined as all remaining genomic regions after excluding CDSs, UTRs, and their corresponding antisense regions. The same approach was applied to the *Eco* MG1655 genome (NC000913.3), with TSS and TTS annotations obtained from SEnd-seq^18^.

### Quantification of pausing frequency and sequence composition bias

For each coding sequence (CDS) and corresponding antisense, transcriptional pausing frequency was quantified as the number of identified pausing sites normalized by gene length. Pausing frequency was the number of pausing sites per kilobase of sequence. To characterize sequence composition, nucleotide frequencies (A, G, C, T) were calculated for each CDS and its antisense sequence. A purine–pyrimidine composition bias was defined as:

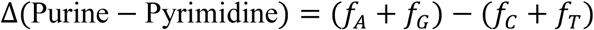

where 𝑓*_NT_* denotes the fractional abundance of nucleotide *N* within the sequence. To assess the relationship between transcriptional pausing and sequence composition, pausing frequency was analyzed as a function of Δ(Purine − Pyrimidine)for CDSs and antisense CDSs. Genes were grouped into bins based on their purine–pyrimidine bias, and the mean pausing frequency within each bin was calculated. Error estimates were computed as the standard error of the mean (SEM). This analysis was performed independently for *Cdf* and *Eco* RNA polymerase datasets.

## DATA AVAILABILITY

RNAP-seq datasets from this study have been deposited in the Gene Expression Omnibus (GEO) with the accession number GSE329416.

## CODE AVAILABILITY

The custom scripts used in this study are available on Zenodo (https://zenodo.org/records/19837913). Other data that support the findings of this study are available from the corresponding author upon request.

